# Integrative pipeline to profile and target endocrine therapy-insensitive cell populations in ER+ breast cancer

**DOI:** 10.64898/2026.09.10.750719

**Authors:** Svetlana E. Semina, Rosemary J. Huggins, Huiping Zhao, Keiko Yanagihara, Monica Sheinin, Virgilia Macias, Fatimah Alani, Leonid Feferman, Andre A. Kajdacsy-Balla, Debra A. Tonetti, Kent F. Hoskins, Geoffrey L. Greene, Jonna Frasor, Jonathan L. Coloff

## Abstract

Up to 40% of patients with estrogen receptor positive breast cancer will experience relapse, either while on endocrine therapy (ET) or after ET is completed. A major contributor to ET failure is the presence of ET-insensitive cell populations within tumors. Here, we developed an analytical pipeline to systematically identify and target these populations by integrating single-cell RNA sequencing of ER+ tumors from the FELINE clinical trial with functional validation in a panel of patient-derived xenograft organoid models. We found that ET-insensitive cells are detected in all tumors regardless of clinical response and exhibit higher transcriptional heterogeneity than ET-sensitive populations. Using our pipeline, we identified and validated new therapeutic options that target patient-specific and shared ET-insensitive populations. Our integrated workflow provides a robust platform for identifying and targeting ET-insensitive cells and offers a translational framework to develop precision medicine approaches to improve outcome in breast cancer patients.

## Introduction

Despite the success of endocrine therapy (ET) and the development of new second-line therapies such as CDK4/6 inhibitors, up to 40% of patients with estrogen receptor positive (ER+) breast cancer will relapse(1–4). A major contributor to ET failure is tumor heterogeneity, which exists at both intertumoral and intratumoral levels. Intertumoral heterogeneity reflects differences in ET response between patients, while intratumoral heterogeneity refers to the coexistence of distinct cell populations within a single tumor, each with varying sensitivity to the therapy.

Multiple studies have been devoted to breast tumor heterogeneity, aiming to understand how it affects ET sensitivity. For example, one study demonstrated the evolution of malignant cell populations over years of treatment, revealing pre-existing and acquired drug-resistant phenotypes using scRNA-seq(5). Another group developed an *ex vivo* operating room-to-laboratory pipeline and identified several ET-responsive and non-responsive cell populations in human specimens(6). A more integrative analysis has also addressed both inter- and intratumoral heterogeneity in attempts to uncover mechanisms of therapy resistance(7). These examples highlight recent advances in unraveling tumor heterogeneity and characterizing ET-insensitive cell populations. However, it is important to note that these studies are limited by the number and type of samples.

In addition to tumor heterogeneity, the treatment of patients with ER+ tumors is further complicated by the varied expression and activity of ER itself(8). While ER status is a key determinant of ET eligibility, it does not fully capture the functional diversity of ER signaling at the single-cell level. Tumors classified as ER+ often harbor subpopulations with low or absent ER activity, which may evade therapy and contribute to disease progression(9). Historically, studies of ET resistance have relied heavily on established cell lines such as MCF7 and T47D that are highly enriched with ER+ cells(10). Although informative, these models lack the molecular and cellular diversity of patient tumors and may not accurately reflect the dynamics of ET response in the clinical setting.

To address these limitations, we present an integrated framework that combines clinical and experimental data to systematically identify and target ET-insensitive cell populations. Leveraging scRNA-seq data from the FELINE clinical trial, we profiled tumors from ER+ breast cancer patients treated with letrozole at baseline and after 14 days of therapy(11–13). We found that ET-sensitive populations are relatively conserved across patients, whereas ET-insensitive populations are more heterogeneous and patient-specific. To functionally validate these findings and target ET-insensitive cell populations, we built a panel of patient-derived xenograft organoid (PDxO) models that retain key features of the original tumors including the presence of heterogeneous ET-insensitive populations. These models enabled us to systematically identify and target these populations using our predictive therapeutic pipeline that integrates single-cell clustering, differential gene expression analysis, and drug response profiling. This approach allowed us to uncover both patient-specific and shared ET-insensitive cell populations, and to test tailored therapeutic strategies for targeting ET resistance. Together, our findings reveal a complex and heterogeneous landscape of endocrine therapy response, highlighting the critical importance of single-cell resolution to inform therapeutic strategies aimed at overcoming resistance and improving patient outcome.

## Results

### ET-insensitive cell populations are present in all tumors irrespective of clinical response

To investigate the cellular basis of ET response in clinical samples of ER+ breast cancer, we analyzed scRNA-seq data from nine tumors collected during the FELINE clinical trial(11). Tumor samples were obtained at baseline (Day 0) and after 14 days of letrozole treatment - a schedule that closely mirrors our previously established *in vitro* protocol for identifying ET-insensitive cells(13). Based on clinical outcomes, five tumors were classified as responsive and four as non-responsive. For each patient, cells from Day 0 and Day 14 were integrated in an individual dataset and clustered using the Seurat package(14), revealing multiple transcriptionally distinct clusters in each tumor (Fig. 1 A-B). To determine which clusters were ET-insensitive, we performed differential abundance analysis where clusters statistically significantly depleted after treatment were classified as ET-sensitive, while those enriched or unchanged were considered to be ET-insensitive(13). This analysis revealed that all tumors - regardless of clinical classification - contain both ET-sensitive and ET-insensitive cell populations (Fig. 1 C, S. Table 1).

**Figure 1.**
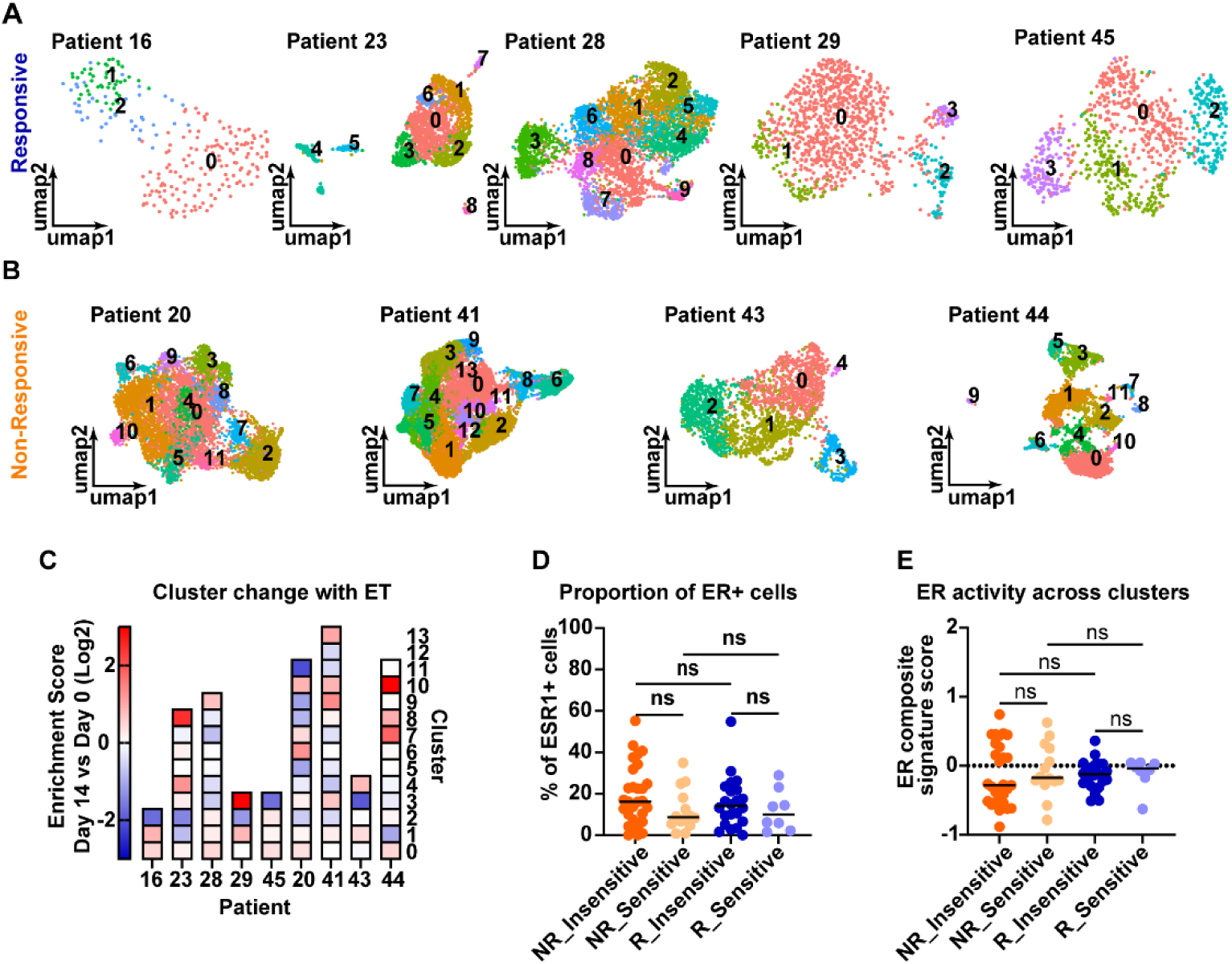
ET-insensitive cell populations are present in all ER+ tumors regardless of clinical response. **(A – B)** UMAP plots of integrated single-cell RNA-seq datasets from nine ER+ breast tumors collected at Day 0 and Day 14 of letrozole treatment in the FELINE clinical trial. Each tumor was analyzed independently. **(C)** Heatmap summarizes the results of patient-specific differential abundance analysis for tumors from (A) and (B) across all clusters. Enrichment scores (Log2) and associated p-values are provided in S. Table 1. Cluster identities are unique to each patient and are not intended for direct comparison across tumors. **(D)** Distribution plot of ER+ cells proportions in each cluster. NR = Non-Responsive, R = Responsive. **(E)** Distribution plot of composite ER scores (mean of ER-associated signature scores per cluster) grouped by clinical outcome and ET sensitivity. Each point represents a cluster. For (D, E) statistical testing was performed using one-way ANOVA with multiple comparisons, ns = not significant.

To further characterize these populations and investigate whether ER expression or signaling activity underlies differential ET response, we examined proportion of ER+ cells and ER pathway activity across clusters in each patient. We identified clusters enriched for ER+ cells and signaling, as well as clusters with low or absent ER expression/activity, in all tumors irrespective of their clinical classification (S. Table 2) (15). Moreover, statistical comparisons of group means revealed no significant differences in the proportion of ER+ cells or ER signaling activity between ET-sensitive and ET-insensitive clusters (Fig. 1 D–E). In parallel, linear regression models applied to each individual ER-related signature and the composite ER score similarly showed no significant associations (S. Table 3). These findings indicate that neither ER expression nor ER signaling activity consistently distinguish ET-sensitive from ET-insensitive clusters. Additionally, to account for potential bias due to unequal cell numbers across samples, we performed patient-level analysis by summarizing ER+ proportions and ER activity per tumor and clinical response. Consistent with cluster-level results, no significant differences were observed between responsive and non-responsive patients (S. Fig. 1 A–D, S. Table 4).

These findings suggest that ET-insensitive cell populations are an intrinsic feature of ER+ breast tumors and are present regardless of clinical outcome. Furthermore, while ER expression and activity are important for clinical treatment decisions, they may not fully capture the complexity of ET sensitivity, suggesting that additional transcriptional programs may be predictive of endocrine resistance.

### ET-insensitive populations are more transcriptionally diverse than ET-sensitive populations

We next asked whether ET-sensitive and ET-insensitive cell populations are common across patients. To address this, we extracted ET-sensitive and ET-insensitive cell populations from all tumors and re-clustered them into two integrated datasets. Interestingly, clusters derived from ET-sensitive cells are broadly distributed across patients (Fig. 2 A–D, S. Table 5), suggesting that these populations are more conserved and shared. In contrast, ET-insensitive cells form fifteen distinct clusters with variable prevalence across patients (Fig. 2 E–H, S. Table 6), demonstrating greater inter- and intrapatient heterogeneity. To quantify this heterogeneity, we used Simpson’s diversity index. ET-sensitive clusters generally exhibited higher Simpson’s diversity scores, consistent with these populations being more broadly shared across patients, while ET-insensitive clusters exhibited a broader range of Simpson’s diversity scores. Five ET-insensitive clusters (Clusters 1, 6, 7, 12, and 14) were patient-specific, whereas the remaining clusters were represented across several or all patients (Fig. 2I).

**Figure 2.**
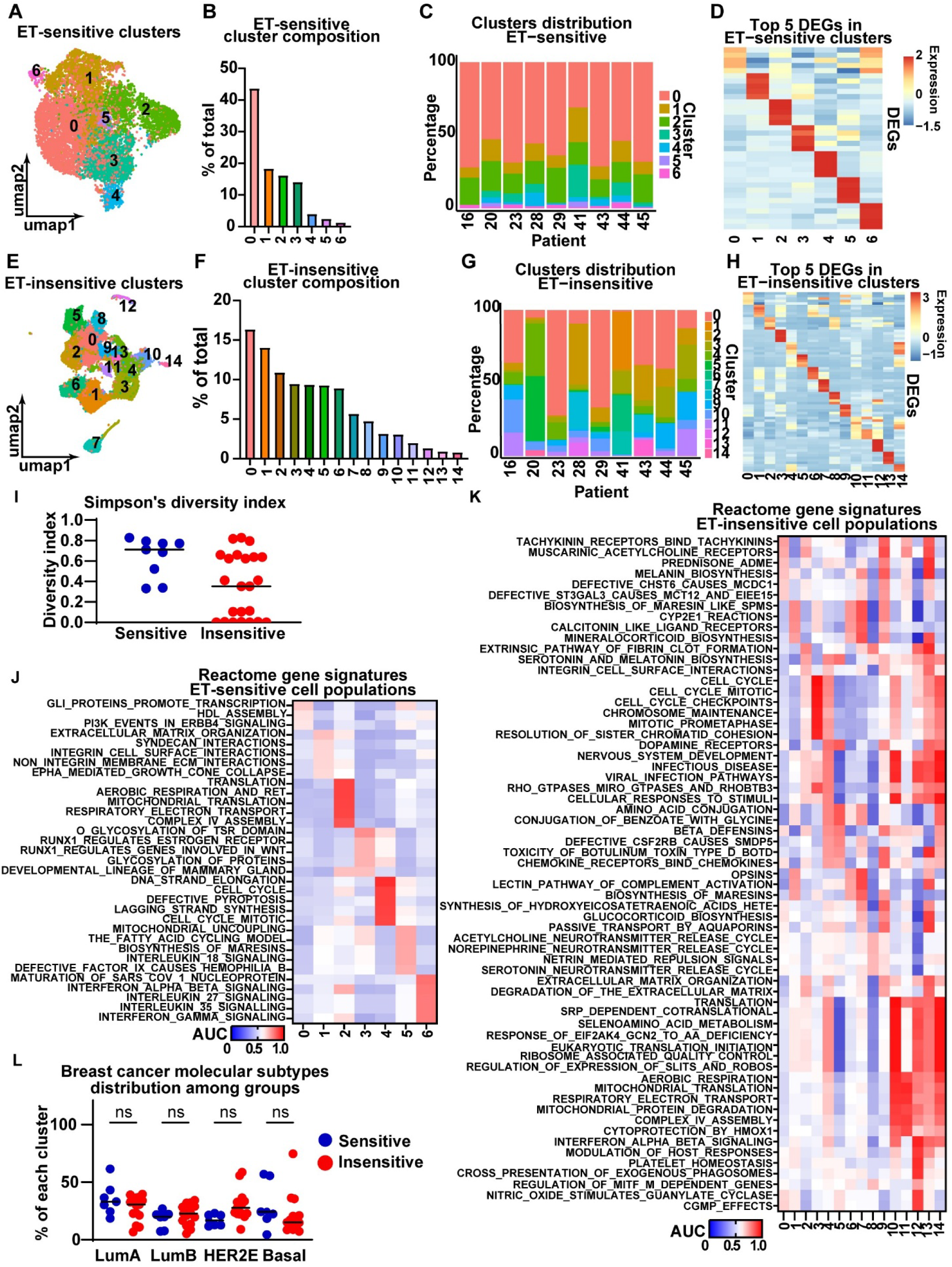
ET-insensitive populations are more heterogeneous and transcriptionally diverse than ET-sensitive populations. **(A-D)** Characterization of ET-sensitive populations: **(A)** Integrated UMAP of ET-sensitive cells from Fig. 1A-B; **(B)** Relative abundance of ET-sensitive clusters; **(C)** Distribution of ET-sensitive clusters across patients; **(D)** Top 5 up-regulated and differentially expressed genes in each cluster within ET-sensitive cell populations. **(E-H)** Characterization of ET-insensitive populations: **(E)** Integrated UMAP of ET-insensitive cells from Fig. 1A-B; **(F)** Relative abundance of ET-insensitive clusters; **(G)** Distribution of ET-insensitive clusters across patients; **(H)** Top 5 up-regulated and differentially expressed genes in each cluster within ET-insensitive cell populations. **(I)** Simpson’s diversity index for all ET-sensitive and ET-insensitive cell population is shown in box plot. The lines indicate mean values across the clusters in each group, each dot represents one cluster. **(J, K)** FEA of Reactome gene signatures was performed on ET-sensitive **(J)** and ET-insensitive **(K)** clusters. AUC values are shown in a heatmap, and p-values are presented in S. Tables 7 and 8. **(L)** Distribution plot shows proportion of cells assigned to each molecular subtype (LumA, LumB, HER2E, Basal) across ET-sensitive and ET-insensitive clusters. Mean subtype proportions were compared between groups using one-way ANOVA with multiple comparisons, ns = non-significant. Each point represents a cluster.

To assess transcriptional heterogeneity in both groups, we performed functional enrichment analysis (FEA) using Reactome gene signatures. ET-sensitive clusters are enriched for pathways primarily reflecting cell cycle–associated processes, respiration, inflammation, cell membrane and extracellular matrix organization (Fig. 2 J, S. Table 7). In contrast, ET-insensitive clusters exhibit markedly heterogeneous enrichment profiles, with variable activation of diverse biological processes, including proliferation, metabolism, cellular stress response, protein processing, and signal transduction (Fig. 2 K, S. Table 8), suggesting that ET-insensitive cell populations use different mechanisms to overcome selective pressure of ET. Together, these findings demonstrate that ET-insensitive cell populations are more heterogeneous and transcriptionally diverse than ET-sensitive populations. Notably, all four major molecular subtypes - luminal A, luminal B, HER2-enriched, and basal (16) - are present in both ET-sensitive and ET-insensitive clusters, with no significant differences in their proportions (Fig. 2 L). Additionally, no significant differences in subtype composition are observed between responsive and non-responsive patients or ET-sensitive and ET-insensitive cells within each tumor (S. Fig. 2). This suggests that ET sensitivity is not strictly defined by classical molecular subtype, but rather by the transcriptional and functional state of individual cell populations.

### ET-insensitive gene signatures are predictive of poor patient outcome and worse clinical-pathological features

To assess whether ET-insensitive populations, despite their patient-specific nature, harbor transcriptional programs linked to poor clinical outcomes, we generated custom gene signatures based on the top 200 differentially expressed genes (DEGs) from each cluster identified in Fig. 2 E (S. Table 9) and applied these signatures to untreated ER+ breast tumors from the METABRIC bulk gene expression dataset(16,17). Although single-cell derived signatures represent transcriptional programs of distinct cell populations, we tested their detectability in bulk RNA-seq data to evaluate whether these populations are transcriptionally dominant to influence bulk-level measurements. Among the ET-insensitive clusters, signatures from three (Clusters 2, 3, and 9) are significantly associated with worse overall survival (Fig. 3 A) and increased risk of relapse (Fig. 3 B, S. Table 10). Several signatures also correlate with adverse clinical features, including more aggressive molecular subtypes and higher tumor cellularity (Fig. 3 C, S. Table 11). Surprisingly, only two signatures from ET-sensitive clusters (Clusters 0 and 5) are significantly associated with favorable outcome (S. Table 10). To identify key contributing genes within cluster-derived signatures, we performed LASSO regression analysis, which prioritized genes with the strongest contribution to each model (S. Tables 12–13). Signatures from Clusters 2, 3, and 9 were distinct and varied by clinical endpoint, such as relapse-free survival (RFS) and overall survival (OS). Cluster 3 was enriched for proliferation-associated genes, Cluster 2 reflected diverse programs including proliferation, migration, metabolism, and epigenetic regulation, and Cluster 9 was dominated by mitochondrial and metabolic genes. A descriptive overlap analysis showed varying degrees of overlap between the LASSO derived RFS- and OS-associated prognostic gene sets across clusters, with greater overlap observed for Cluster 3 than for other clusters (Fig. 3 D). Together, these findings suggest that multiple ET-insensitive transcriptional programs are independently associated with poor clinical outcomes while retaining distinct prognostic gene compositions.

**Figure 3.**
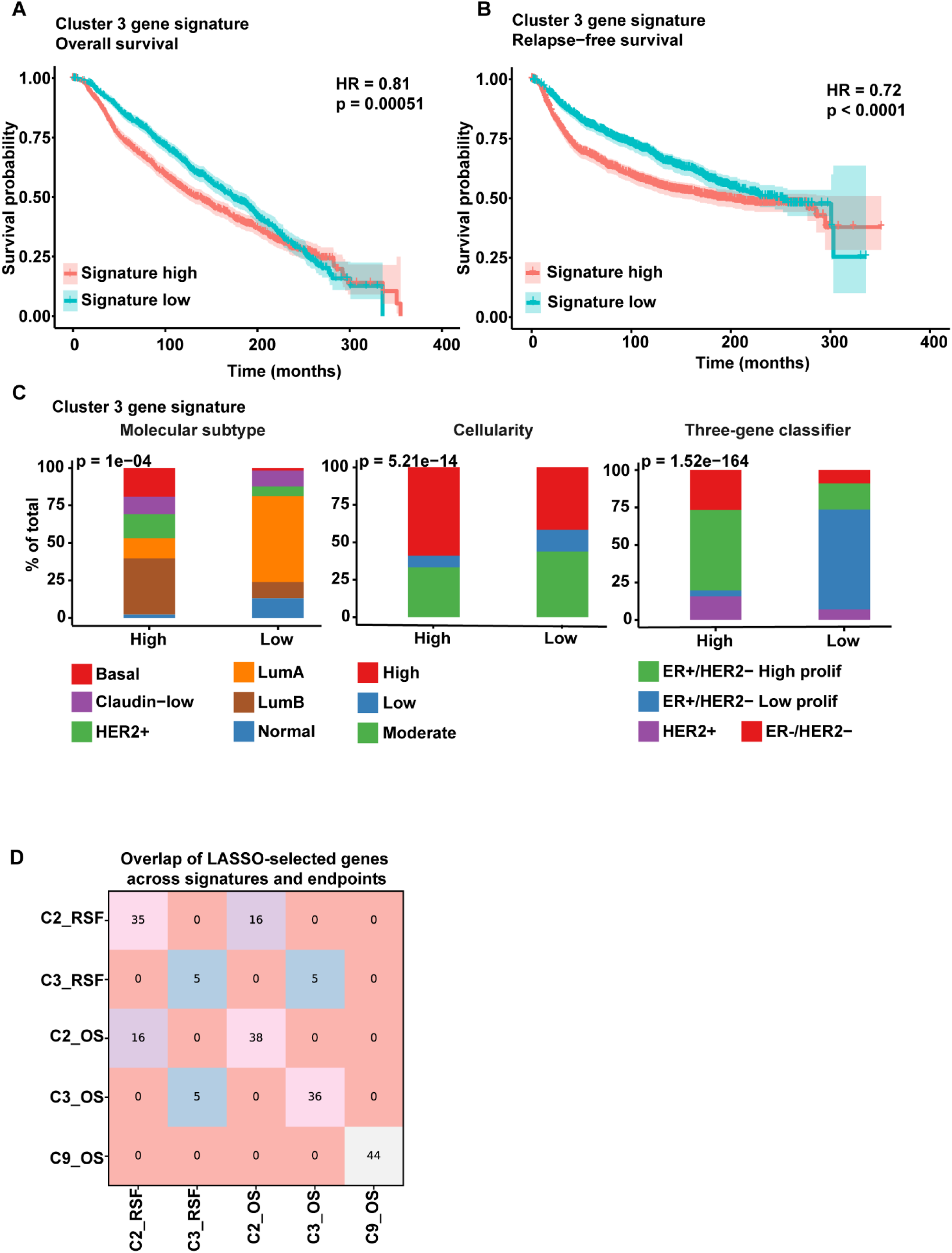
Gene signatures derived from ET-insensitive clusters are predictive of poor clinical outcome. **(A–C)** Gene signatures from ET-insensitive and -sensitive clusters were interrogated in 1817 ER+ breast tumors from METABRIC cohort available through cBioPortal. Patient overall **(A)**, relapse-free survival **(B)**, and clinical pathological features **(C)** were compared between tumors high vs. low for expression of the tested signatures. The results for ET-insensitive Cluster 3 gene signature are displayed. Statistical significance was determined using log-rank test (A-B) or chi-square test (C). Full statistical analyses for all signatures are presented in S. Tables 10 and 11. **(D)** Overlap of genes selected by LASSO regression for signatures derived from ET-insensitive clusters (Fig. 2E) across recurrence-free survival (RFS) and overall survival (OS). Values indicate the number of shared genes between signature pairs.

### New patient-derived models retain heterogeneity of ER+ breast tumors

The complexity of the ET-insensitive cell populations from patient samples prompted us to build a panel of ER+ breast cancer patient-derived models that recapitulate tumor heterogeneity and allow the study of ET sensitivity in greater detail (S. Table 14). This panel includes four organoid models derived from patient-derived xenografts (PDxOs). Two models, HCI003 and HCI017, are ER+/PR+ and were derived from treatment-naïve primary tumors (generously provided by Dr. Welm (18)), while two additional models, UIC013 and UIC020, were developed at UIC and the University of Chicago. UIC013 was derived from a treatment-naïve ER+/PR- tumor, and UIC020 originated from a tumor that initially presented as ER- but recurred as ER+. Since UIC013 and UIC020 are newly developed models, we first sought to confirm that both the PDXs and PDxOs faithfully preserve key features of the original patient tumors.

To assess molecular fidelity, we first performed NanoString Breast Cancer 360 biological signatures analysis on primary tumors and matched PDXs for UIC013 and UIC020 (S. Fig. 3 A-D). As expected, immune and inflammatory signatures are markedly reduced in PDXs due to the use of immunodeficient mice. Notably, both tumors and PDXs are enriched for basal-like or basal/HER2 molecular subtype on their global expression profiles. However, ER expression, which was also confirmed by IHC (S. Fig. 3 E, F), and ER signaling pathways are still detectable, suggesting that these tumors display ER-low phenotypes. Most other molecular signaling pathways, including AR, FOXA1, and hypoxia, are preserved in PDX models.

We next evaluated whether the cellular heterogeneity of the original tumors was preserved in both PDXs and developed PDxOs. Spatial transcriptomics was performed on FFPE-preserved primary and PDX tumors, and single-cell RNA sequencing was conducted on the corresponding PDxOs (Fig. 4 A-C, S. Fig. 4 A-C). Each dataset was analyzed independently using the Seurat package. To identify epithelial-enriched spots in patient tumors and PDXs, spatial transcriptomics data were annotated using a single-cell breast cancer atlas(19) (S. Fig. 4 D). Epithelial-enriched spots from the primary tumors were clustered separately for UIC020 (Fig. 4 D-E) and UIC013 (S. Fig. 4 E-F), and the distribution of these clusters was then assessed in the matched PDX and PDxO datasets using Seurat’s label-transfer method. This analysis revealed that all epithelial subpopulations present in the original tumors are detected in the matched PDX and PDxO models, with some variation in prevalence (Fig. 4 F, S. Fig. 4 G).

**Figure 4.**
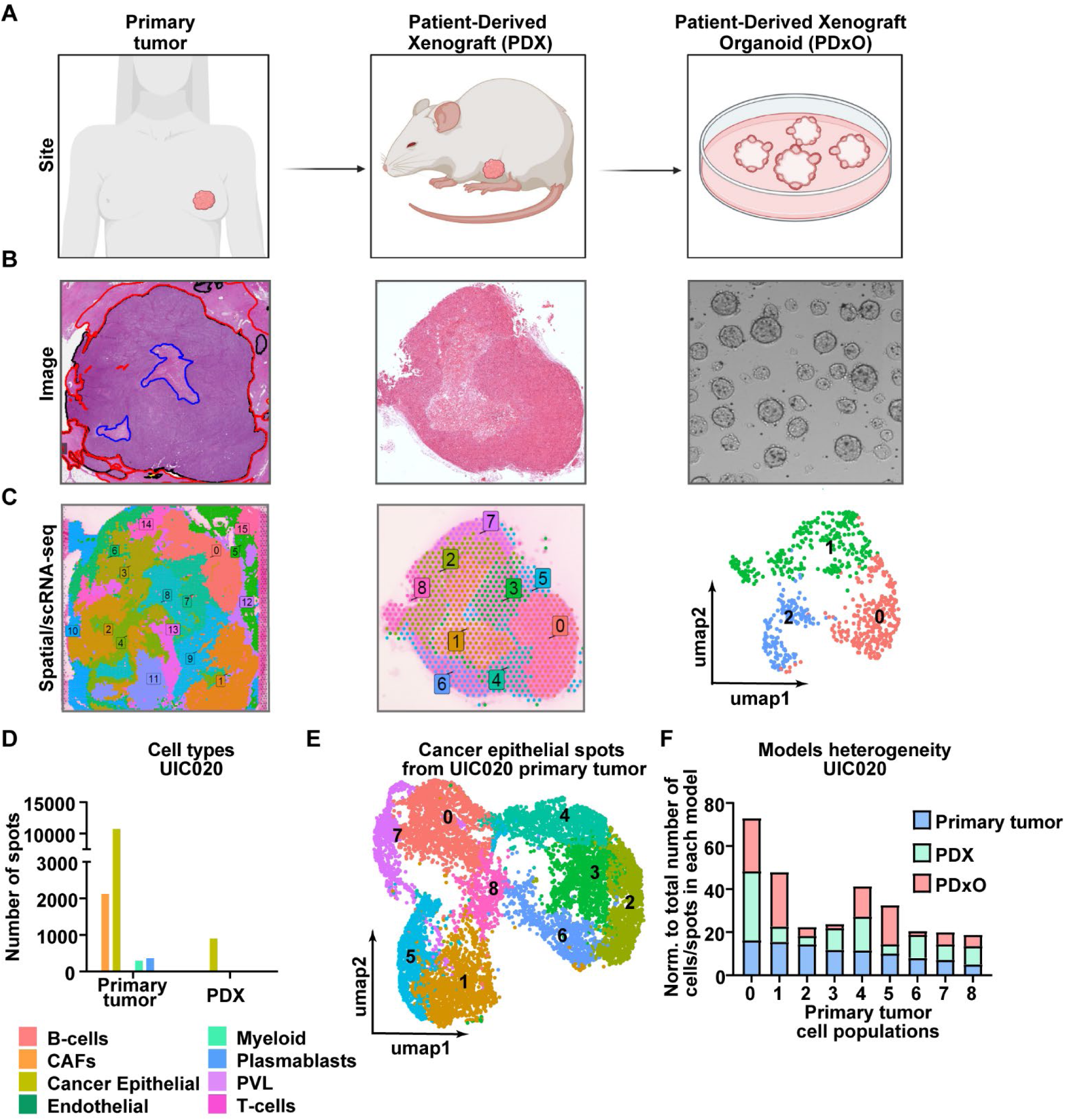
Newly developed UIC020 PDxO model retains molecular and cellular heterogeneity of the original ER+ breast tumor. **(A–C)** Schematic representation and characterization of matched primary tumor, patient-derived xenograft (PDX), and patient-derived organoid (PDxO) models from UIC020. **(A)** Schematic overview of models’ development. **(B)** Visual representation of each UIC020 model. Hematoxylin and eosin (H&E) staining was used to visualize primary and PDX tumors, and brightfield microscopy was used to visualize PDxOs. Morphological annotations of the primary tumor regions are labeled: black - tumor, red - fibrosis, blue - necrosis. **(C)** Spatial clustering of primary tumor and PDX samples aligned to H&E-stained sections. UMAP plots show clustering of PDxO cells. **(D) (E)** Bar plot displays cell type distribution across spatial RNA-seq spots from UIC020 primary tumor and PDX. **(F)** The UMAP plot shows clustering of cancer epithelial spots from UIC020 primary tumor. **(G)** Bar plot shows the distribution of epithelial clusters from the UIC020 primary tumor across PDX and PDxO models. Label transfer analysis was used to assess conservation of epithelial subpopulations across models. Figure 4A was created in BioRender. Semina, S. (https://BioRender.com/8acdtax) is licensed under CC BY 4.0.

Given that loss of ER expression and functionality is a common challenge in ER+ PDxO development, we next validated ER across all models. We confirmed ER expression by immunofluorescence (IF) for ERα protein (S. Fig. 5 A, B), and the proportion of ER+ cells classified in our models as ER-low to moderate. To confirm ER functionality, we evaluated ET sensitivity in all four PDxO models using a 14-day treatment with fulvestrant (ICI-182,780), 4-hydroxytamoxifen (4OHT), or vehicle control (Veh). Previously published data for HCI003 and HCI017 (13) were extended with new assays for UIC013 and UIC020. All models show reduced organoid area after ET, even after 20 weeks of passaging, indicating partial responsiveness and functional ER signaling (S. Fig. 5 C, D). The decrease in S-phase proliferative cells following ET treatment was also confirmed by EdU assays, even when changes in organoid area were not statistically significant (S. Fig. 5 E-H), likely reflecting heterogeneity in the response to ET.

**Figure 5.**
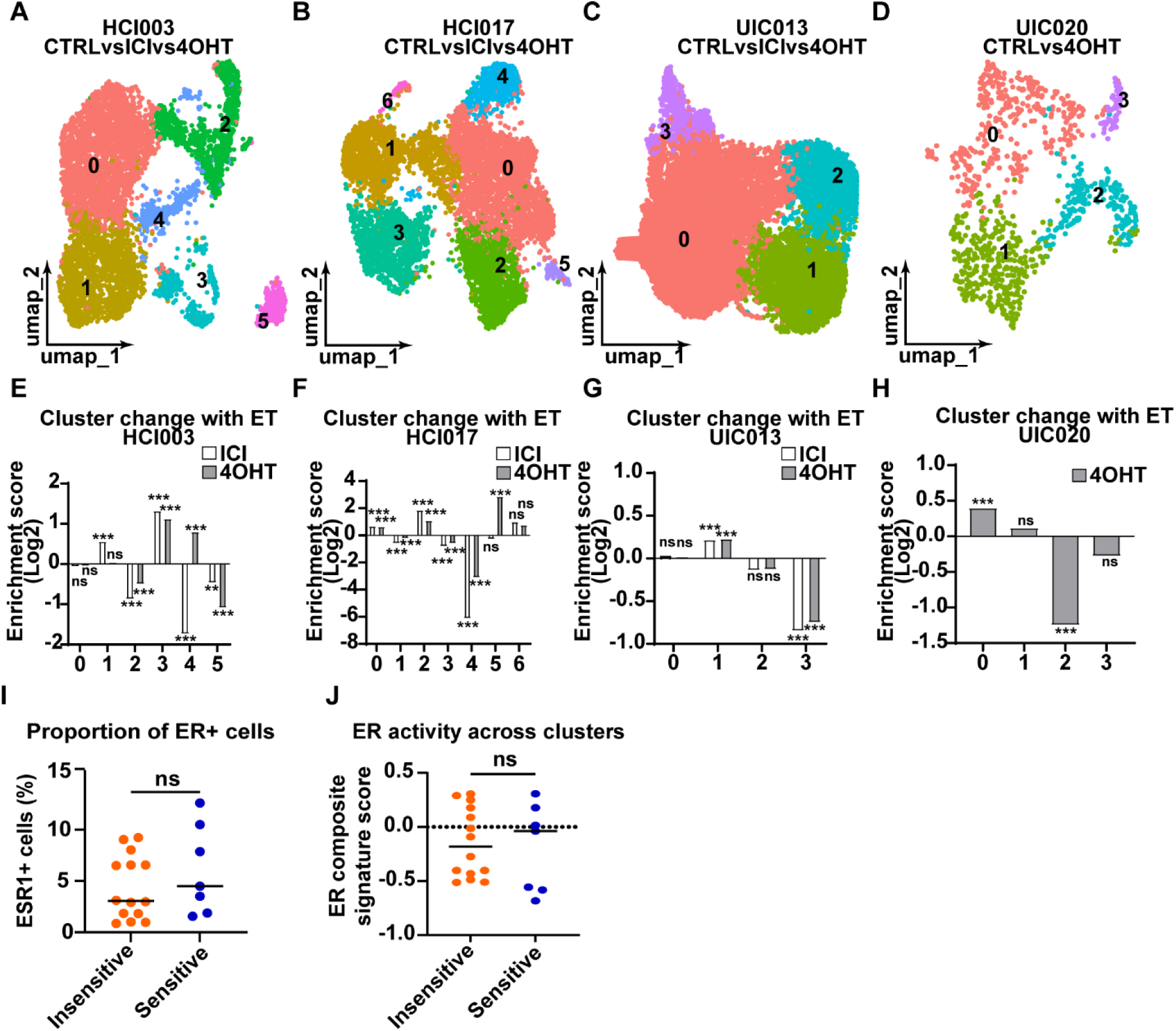
ET-insensitive populations are present in PDxO models and mirror clinical observations. **(A–D)** UMAP plots and integrated clustering of scRNA-seq data from each PDxO model treated with 1uM of ICI, 4OHT, or vehicle (control) for 14 days. **(E–H)** Differential abundance analysis shows enrichment score and ET-sensitive clusters (statistically significantly depleted post-treatment) or ET-insensitive clusters (enriched or unchanged post treatment) for each model. **P < 0.01, ***P < 0.001, ns = not significant. **(I)** ESR1+ cell proportions across clusters and ET sensitivity groups. **(I)** Distribution plot shows proportions of ER+ cell in each cluster grouped by ET sensitivity. **(J)** Distribution plot shows composite ER scores (mean of ER-associated signature scores per cluster) grouped by ET sensitivity. Each point represents a cluster. For (I, J) statistical testing was performed using one-way ANOVA, ns = not significant.

We next performed scRNA-seq on the PDxO models under standard growth conditions (19) to characterize cellular heterogeneity and ER signaling at single-cell resolution (S. Fig 6 A-D). We first assessed the proportion of ER⁺ cells in each model and found that it was lower (S. Fig 6 E) than that observed by immunofluorescence (S. Fig.5 A, B), consistent with the “drop-out” effect commonly observed in scRNA-seq studies. Unsupervised clustering nevertheless identified distinct cell populations within each model, with variable ER expression and activity across clusters (S. Fig. 6 F, G, S. Table 15). These findings demonstrate that our PDxO panel preserves the molecular and cellular heterogeneity of ER+ breast tumors, retains ER expression and activity, and remains responsive to ET. These models provide a robust platform for further investigation of ET sensitivity and new therapeutic options.

**Figure 6.**
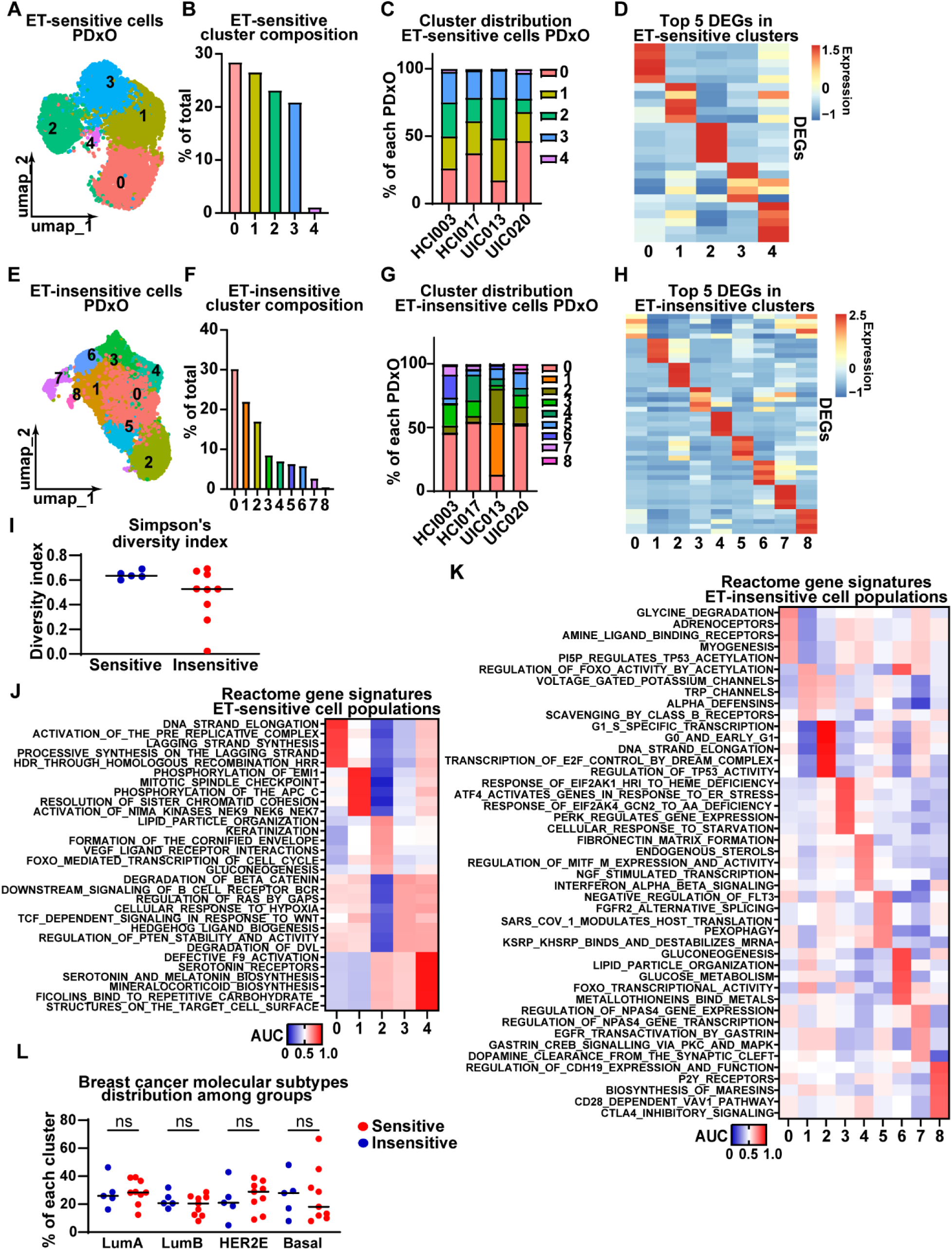
ET-insensitive populations in PDxOs are transcriptionally diverse and patient-specific. **(A-D)** Characterization of ET-sensitive populations: **(A)** Integrated UMAP of ET-sensitive cells from Fig. 5A-D; **(B)** Relative abundance of ET-sensitive clusters; **(C)** Distribution of ET-sensitive clusters across patients; **(D)** Top 5 up-regulated and differentially expressed genes in each cluster within ET-sensitive cell populations. **(E-H)** Characterization of ET-insensitive populations: **(E)** Integrated UMAP of ET-insensitive cells from Fig. 5A-D; **(F)** Relative abundance of ET-insensitive clusters; **(G)** Distribution of ET-insensitive clusters across patients; **(H)** Top 5 up-regulated and differentially expressed genes in each cluster within ET-insensitive cell populations. **(I)** Simpson’s diversity index for all ET-sensitive and ET-insensitive cell population is shown in box plot. The lines indicate mean values across the clusters in each group. **(J, K)** FEA of Reactome gene signatures was performed on ET-sensitive **(J)** and ET-insensitive (**K)** clusters. AUC values are shown in a heatmap, and p-values are presented in S. Tables 20 and 21. **(L)** Distribution plot shows proportion of cells assigned to each molecular subtype (LumA, LumB, HER2E, Basal) across ET-sensitive and ET-insensitive clusters. Mean subtype proportions were compared between groups using one-way ANOVA with multiple comparisons, ns = non-significant. Each point represents a cluster.

### ET-insensitive cell populations in PDxOs mirror clinical datasets

To experimentally identify ET-insensitive cell populations in our PDxO panel, we mirrored the experimental and analytical workflow used for the FELINE clinical dataset. We performed scRNA-seq on PDxO models treated for 14 days with ICI, 4OHT, or vehicle control (CTRL). Following the same analytical pipeline as in Figures 1 and 2, we conducted unsupervised clustering for each treatment condition within each PDxO model. ET-sensitive and -insensitive cell populations were then defined using differential abundance analysis (Fig. 5 A-H, S. Table 16). Consistent with our findings in the FELINE dataset, each PDxO model contains both ET-sensitive and ET-insensitive cell populations. Furthermore, the proportion of ER+ cells and ER signaling activity within each cluster are not statistically significantly associated with ET sensitivity at the population level, despite the observed trend toward higher ER activity and ESR1+ cell proportions in ET-sensitive populations (Fig. 5 I, J, S. Tables 17,18).

Next, we extracted ET-sensitive and ET-insensitive cells from all PDxO models and re-clustered them into integrated datasets. ET-sensitive cells form five clusters shared across all models, indicating a conserved transcriptional program. While ET-insensitive cells consist of nine distinct clusters with variable prevalence across models, reflecting a greater degree of heterogeneity and revealing both shared and patient-specific populations (S. Table 19). Some clusters, such as Clusters 0, 2, and 5, are shared across four models, while others are enriched in only a subset of models (e.g., Cluster 4) or are highly enriched in individual patients (e.g., Cluster 1). This was further supported by Simpson’s diversity index, which confirmed that ET-insensitive populations are more model-specific and heterogeneous compared to ET-sensitive populations (Fig. 6 A-I).

We next assessed transcriptional heterogeneity using FEA of Reactome gene signatures. Consistent with findings from the FELINE clinical dataset, ET-sensitive clusters derived from PDxO models show similar patterns of pathway enrichment. Specifically, three clusters are enriched for pathways primarily related to proliferation, ER-associated transcriptional response, and metabolic programs (Fig. 6 J, S. Table 20). In contrast, ET-insensitive clusters from PDxO exhibit diverse transcriptional profiles, including pathways associated with stress response, inflammation, and transcriptional regression, together forming more complex pathway activation patterns and indicating greater transcriptional heterogeneity (Fig. 6 K, S. Table 21). Notably, both ET-sensitive and ET-insensitive clusters include cells from all four major molecular subtypes without statistically significant differences in their proportions between groups (Fig. 6 L), further supporting that ET sensitivity is not strictly defined by classical subtypes. These findings demonstrate that the PDxO models recapitulate key features of the ET response observed in patient tumors.

### PDxO ET-insensitive cell populations are detected in patient tumors and are predictive of poor patient outcome

Given the alignment between findings in our PDxO models and the FELINE clinical trial, we used custom gene signatures derived from the top 200 DEGs for each cluster identified in Fig. 6E (S. Table 22) and investigated whether these populations can be detected in untreated human tumors and are predictive of patient outcome. We interrogated nine treatment naïve ER+ breast tumors from the Breast Cancer Atlas(19) (S. Fig. 4D) and eight ET-treated breast tumors from FELINE clinical trial (Fig. 1) and estimated percent of cells harboring similar transcriptional profiles with PDxO ET-insensitive cell populations (Fig. 7 A-C). Across both datasets, we observed highly variable proportions of cells whose transcriptional profiles closely matched to PDxO ET-insensitive populations, indicating that some transcriptional states are recurrent and detectable in primary tumors.

**Figure 7.**
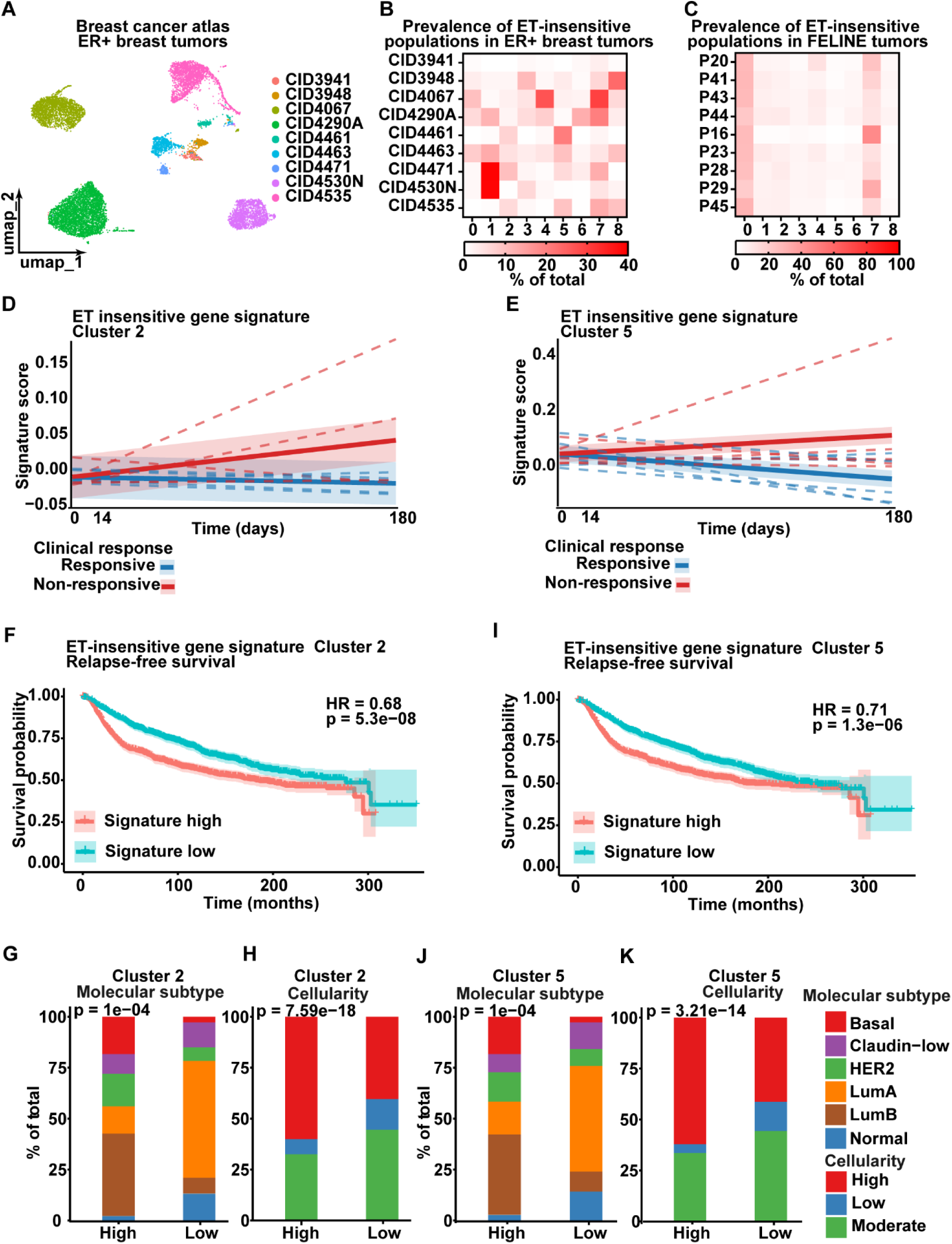
Clinical relevance of ET-insensitive populations derived from PDxOs. **(A)** UMAP plot represents clustering of nine ER+ breast treatment-naïve tumors from breast cancer atlas. **(B)** Prevalence of cells with transcriptional profiles similar to PDxO-derived ET-insensitive populations in ER+ breast tumors (A). **(C)** Prevalence of cells with transcriptional profiles similar to PDxO-derived ET-insensitive populations in ER+ breast tumors from FELINE clinical trial (Fig. 1). **(D-E)** Longitudinal analysis for enrichment of ET-insensitive gene signatures from Cluster 2 and 5 in FELINE tumors (Fig 1) over time. Inter-patient variability in signature activity is shown by dashed lines indicating patient-specific fitted trend estimates and solid lines represent fitted trends from the cell-level mixed-effects model. Interaction estimates and statistical results are reported in S. Table 23. Patient-level sensitivity analyses are reported in S. Table 24. **(F-K)** Gene signatures from ET-insensitive clusters were interrogated in 1817 ER+ breast tumors from METABRIC cohort available through cBioPortal. Patient relapse-free survival **(F,I)** and clinical pathological features **(G,H,J,K)** were compared between tumors high vs. low for expression of the tested signatures. The results for ET-insensitive Cluster 2 and Cluster 5 gene signature are displayed. Statistical significance was determined using log-rank test **(F,I)** or chi-square test **(G,H,J,K)**. Full statistical analyses for all signatures are presented in S. Tables 25 and 26.

To evaluate longitudinal changes during endocrine therapy, we analyzed PDxO-derived ET-insensitive signature activity across all available FELINE timepoints (Day 0, Day 14, and Day 180) using cell-level linear mixed-effects models with patient included as a random effect. Selected signatures derived from Clusters 2 and 5 exhibited significant response-by-time interactions, with non-responsive tumors showing a trend toward higher signature scores at Day 180 in the cell-level analysis (Fig. 7D–E, S. Table 23). To assess the robustness of these findings, we additionally performed a patient-level sensitivity analysis by averaging signature scores within each patient-timepoint sample (S. Table 24). Although the estimated mean signature scores for Clusters 2 and 5 showed increases at Day 180 in non-responsive tumors, these analyses did not reach statistical significance, reflecting the limited number of patients, incomplete longitudinal sampling, and/or substantial inter-patient heterogeneity.

To test the predictive power of PDxO ET-insensitive signatures in bulk RNA sequencing of ER+ tumors, we utilized the METABRIC database, as we did for tumors from FELINE trial (Fig. 3). Among the tested gene signatures, only signatures from Clusters 2, 5, and 7 are predictive of worse overall survival (Fig. 7F) and Clusters 2 and 5 increased risk of relapse (Fig. 7I, S. Table 25). Also, several gene signatures are associated with worse clinical pathological features (Fig. 7 G-K) (S. Table 26). As in the FELINE analysis, we performed LASSO regression on PDxO-derived gene signatures to identify key contributing genes (S. Tables 27, 28). Consistent with the FELINE dataset, we identified one signature dominated by proliferative genes (Cluster 2) and other signatures were characterized by diverse biological programs, including stress response, chromatin regulation, and metabolism (Cluster 5), cytoskeletal organization and vesicle trafficking (Cluster 7). Together, these findings demonstrate that PDxO-derived ET-insensitive transcriptional programs are detectable in patient tumors, are associated with poor clinical outcome and adverse clinicopathological features, and capture biologically distinct transcriptional states with potential clinical relevance.

### Targeting ET-insensitive cell populations with a predictive therapeutic pipeline

To identify therapeutic strategies aimed at eliminating ET-insensitive cell populations, we assembled a predictive therapeutic pipeline that combines various bioinformatic methods and includes following analytical steps: 1) unsupervised clustering of cells treated with vehicle or ET for 14 days, 2) identification of ET-insensitive cell populations, 3) selection of the top 200 DEGs for each ET-insensitive cell population, 4) performing DREEP (DRug Estimation from single-cell Expression Profiles) analysis(20), a computational framework that predicts drug sensitivity in single-cell transcriptional profiles, and 5) testing DREEP predicted drugs in vitro (S. Fig. 7A). DREEP analysis leverages large-scale pharmacogenomic datasets (e.g., Cancer Therapeutics Response Portal v2 - CTRPv2), where gene expressions across cancer cell lines are correlated with drug response to identify gene signatures associated with drug sensitivity and resistance. These signatures are then used to compute drug sensitivity scores (DSS) for each drug at the cell population level, enabling prediction of drug sensitivity and prioritization of candidate compounds targeting transcriptionally distinct populations.

As a proof-of-concept study, we first tested this pipeline on our previously published dataset of 4OHT-treated MCF-7 cells (S. Fig. 7B, C)(13). For in vitro testing, we selected Clusters 1 and 4, as both were enriched following ET treatment. DREEP predicted multiple candidate drugs for each cluster to target, with only twelve compounds overlapping between the two clusters (S. Fig. 7D, E, S. Table 29). We then evaluated the ability of ten of these compounds to target ET-insensitive cell populations in MCF-7 cells using clonogenic assays by treating cells with increasing doses of each compound in the presence or absence of a fixed dose of 4-hydroxytamoxifen (4OHT; 1 μM) for 14 days. Several drugs, including pluripotin and dasatinib, show higher efficacy in reducing ET-tolerant cells (S. Fig. 7F-H). To assess whether these compounds also target cells with established resistance, we performed proliferation assays in three ET-resistant MCF-7-derived cell lines, TAMR (Tamoxifen resistant), FULR (Fulvestrant resistant) and HER2 (HER2 overexpressing) (21,22) (S. Fig. 7I-L). To account for differences in growth rates among the resistant lines, we also performed growth rate correction (S. Fig. 8). These data demonstrate that ET-insensitive cell populations can be systematically identified and targeted using a predictive therapeutic pipeline at the early stages of ET-resistance development to prevent disease progression.

**Figure 8.**
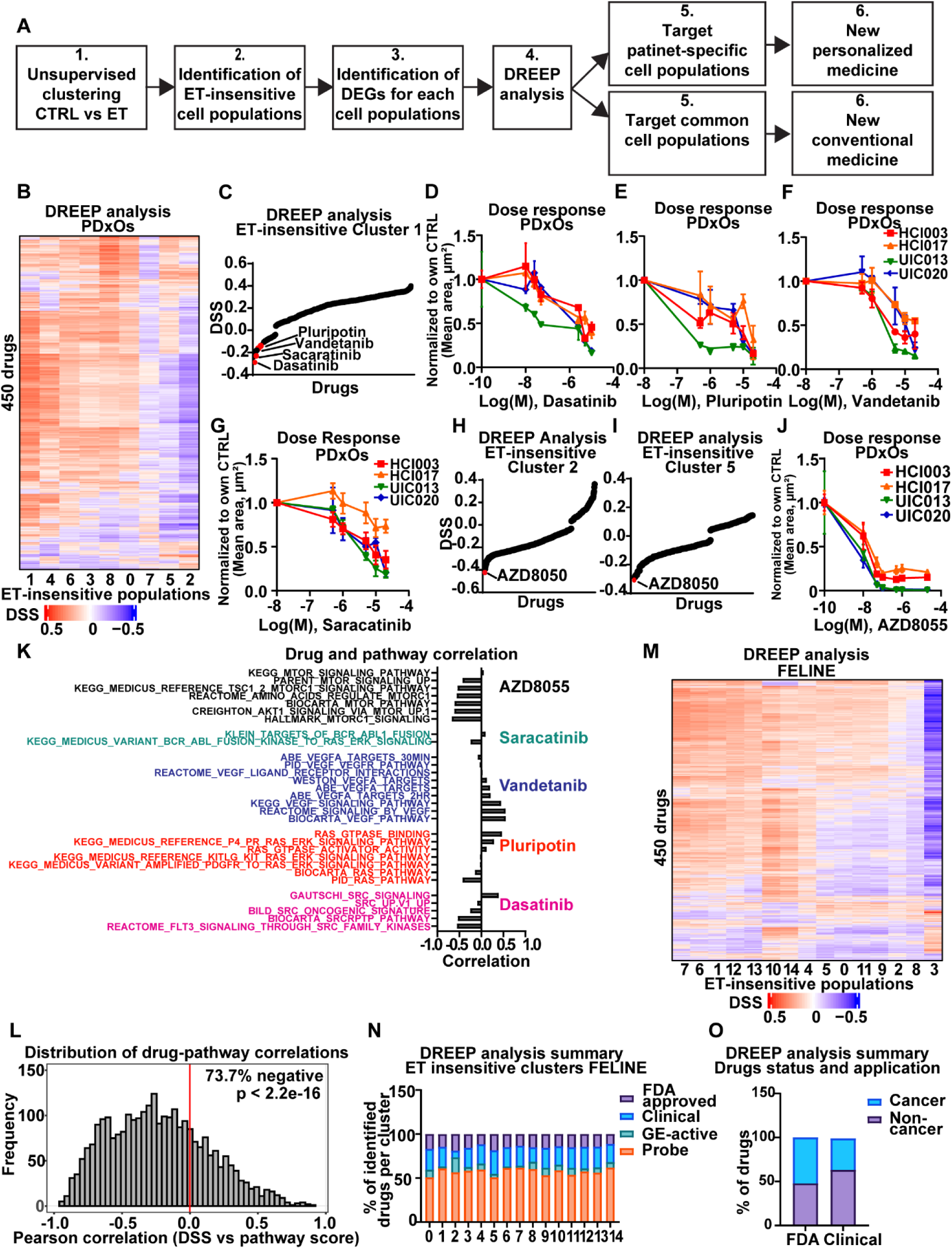
Therapeutic pipeline enables targeting of patient-specific and shared ET-insensitive populations. **(A)** Schematic of the therapeutic pipeline application to PDxO and FELINE datasets. **(B)** Heatmap depicts drug sensitivity scores (DSS) for 450 drugs tested across clusters identified in ET-insensitive cells from PDxO models (Fig. 6E). Negative values indicate sensitivity; positive values indicate resistance. **(C)** DSS for compounds predicted to target ET-insensitive Cluster 1, which is specific to UIC013. **(D-G)** Dose-response assays for drugs predicted to target patient-specific ET-insensitive Cluster 1. **(H,I)** DSS for compounds predicted to target common ET-insensitive Clusters 2 and 5. **(J)** Dose-response to AZD8055 across PDxO models. Full statistical analysis for D-G, J is presented in S. Table 31. **(K)** Bar plot shows correlation between DREEP drug and assisted pathways. **(L)** Distribution of drug-pathway correlations with bi-nominal test results. **(M)** Heatmap depicts DSS for 450 drugs applied to FELINE ET-insensitive clusters (Fig. 2E), as predicted by DREEP analysis. **(N)** Stacked bar plot shows the percentage of predicted compounds per ET-insensitive cluster derived from FELINE clinical trial, stratified by annotation (GE-active = Gene Expression activity). **(O)** Bar plot summarizes the proportion of predicted compounds targeting ET-insensitive clusters from FELINE dataset, stratified by regulatory status (FDA-approved vs. clinical-stage) and cancer relevance.

To test whether our therapeutic pipeline can be used to identify patient-specific therapies to target unique ET-insensitive cell populations, we leveraged our PDxO panel as a clinically relevant platform (Fig. 8A). We first focused on Cluster 1, which is highly specific to UIC013 (Fig. 6G), reasoning that drugs predicted to target this cluster would show higher efficacy in UIC013 compared to other models. DREEP analysis identified several candidate compounds predicted to target the UIC013-specific ET-insensitive Cluster 1, including dasatinib (Src inhibitor), pluripotin (RAS/MAPK inhibitor), vandetanib (VEGFR inhibitor) and saracatinib (Abl1 inhibitor) are more effective at inhibiting growth of UIC013 (Fig. 8B-G, S. Table 30). Dose-response assays followed by replicate-level area under the curve (AUC) analysis demonstrated significantly greater sensitivity of UIC013 to dasatinib and pluripotin compared with the other PDxO models. Vandetanib also showed increased activity in UIC013 relative to HCI017 and UIC020, whereas saracatinib exhibited only modest differences between models (Fig. 8 E-G, S. Table 31). Next, we evaluated Clusters 2 and 5, which are common across multiple PDxO models. AZD8055, an mTOR inhibitor that has been evaluated in an early-phase clinical trial(23), is predicted to target both clusters, and proliferation assays show comparable responses across all models, supporting its potential as a conventional therapy for shared proliferative ET-insensitive populations (Fig. 8H-J).

To assess whether DREEP-predicted drug sensitivities are associated with pathway activity across ET-insensitive clusters, we calculated correlations between pathway gene signature scores from MSigDB and DREEP DSS across clusters for each compound (S. Table 32, 33). As lower DREEP scores indicate higher predicted sensitivity, negative correlations suggest that clusters with higher pathway activity are more sensitive to the corresponding drug. In selected cases, predicted drug sensitivity aligns with pathway activity, including multiple Src-related gene signatures negatively correlating with dasatinib sensitivity (Fig. 8K). However, this relationship was not consistent across all the compounds. This variability likely reflects inherent limitations stemming from differences in the generation of DREEP-derived and pathway gene signatures across diverse models and conditions, as well as the multi-target nature of several compounds. To evaluate this relationship more broadly, we performed a global analysis across all drug-pathway pairs (n = 2,765). This revealed a consistent directional bias toward negative correlations, indicating that higher pathway activity is generally associated with predicted drug sensitivity. Importantly, this trend is observed across multiple pathways rather than being driven by a small subset of drug-pathway pairs, suggesting a system-wide, non-random relationship between transcriptional pathway activity and predicted drug response (Fig. 8L).

Finally, we extended our pipeline to the FELINE clinical dataset, applying DREEP analysis to ET-insensitive clusters identified in patient tumors (Fig. 8M, S. Table 34). We also examined the development status of predicted compounds based on CTRPv2 annotations (e.g., FDA-approved, clinical, preclinical) and assessed their distribution across ET-insensitive clusters (Fig. 8N). Across clusters, approximately 50% of the hits are either FDA-approved or currently under clinical investigation for cancer treatment (Fig. 8O, S. Table 35). Notably, many of these compounds are clinically relevant for breast cancer, including FDA-approved therapies ranging from established agents (e.g., paclitaxel, docetaxel) to more recently approved targeted therapies (e.g., olaparib, alpelisib, neratinib), as well as agents in clinical development (e.g., MK-2206, saracatinib, entinostat). These data suggest that our pipeline can identify therapeutic candidates for a precision oncology approach to treating ET-resistant cell populations that are tailored to specific cell populations.

## Discussion

ET is a central component of treatment for patients with ER+ breast cancer, yet resistance to ET remains a major clinical challenge(6,7). Our study provides a comprehensive analysis of ET response at single-cell resolution, integrating clinical data from the FELINE trial with experimental validation using PDxO models. This approach allowed us to uncover a complex landscape of ET-sensitive and -insensitive cell populations, revealing key insights into the nature of resistance and informing strategies for therapeutic intervention.

Our findings have several important clinical implications. First, the presence of ET-insensitive populations in both responsive and non-responsive tumors suggests that standard ET may suppress sensitive populations while allowing resistant ones to persist or expand. This may explain the low pathological complete response rates observed with neoadjuvant ET, and paradoxical increases in proliferation markers in a subset of luminal A tumors - indicating the presence of treatment-resistant cells and supporting the rationale for expansion of combination therapies that target resistant populations from the outset (24–27).

Second, a central finding of our study is that ET-sensitive cell populations are conserved across patients while ET-insensitive populations are more heterogeneous and patient-specific. This distinction is consistently observed in both clinical samples and PDxO models, suggesting that ET sensitivity is governed by a shared transcriptional program, whereas resistance may arise through diverse molecular mechanisms. While ER expression assessed by IHC remains essential to guide endocrine therapy decisions, our data - alongside previous studies (8,28,29) - show that ER alone has limitations in predicting ET response. These findings highlight the utility of scRNA-seq for a deeper resolution of tumor heterogeneity, uncovering resistant subpopulations that are not detectable by conventional methods. Moreover, transcriptomic analyses capture downstream ER-regulated transcriptional programs but do not directly measure ER protein abundance or post-translational signaling dynamics. Therefore, protein- and transcript-based approaches provide complementary biological information, and their integration may offer a more comprehensive assessment of functional ER signaling, particularly in treatment-resistant tumors. This refinement allows for more precise therapeutic targeting and patient stratification.

Third, our predictive therapeutic pipeline offers a framework for identifying treatments tailored to the transcriptional profiles of resistant populations, with potential applications in precision oncology and expansion to other cancer types. This could allow for drug repurposing to accelerate therapeutic development for patients with ER+ breast cancer, offering a strategic advantage by leveraging existing safety and pharmacokinetic data and enabling faster clinical translation. Importantly, we envision this approach as complementary to existing genomics-based precision oncology strategies, providing functional information on pathway activity and cellular states that may not be fully captured by genomic alterations alone. The observation that some ET-insensitive populations are predicted to respond to inhibitors targeting multiple signaling pathways further highlights the complexity of endocrine resistance and suggests that resistant cell states may depend on interconnected signaling networks rather than a single dominant pathway. Additionally, future studies exploring whether existing therapies, such as CDK4/6 or PI3K inhibitors, can be effective against specific ET-insensitive populations and help identify which patients are most likely to benefit from these agents.

This study has several limitations. While the patient cohort and experimental models used here provide important insight into ET response, they likely do not capture the full spectrum of interpatient heterogeneity in ER+ breast cancer. Additionally, DREEP–pathway associations are not consistent across all compounds, reflecting the complexity of linking transcriptional programs to therapeutic response. Finally, although some prognostic signatures are predictive in bulk RNA-seq datasets, bulk profiling may mask signals from rare but clinically relevant subpopulations.

Our findings build on prior single-cell studies of ET resistance (5) by focusing on early treatment response (Day 0 and Day 14), providing a snapshot of how resistant populations begin to persist under therapeutic pressure before resistance becomes clinically evident. By integrating clinical data with PDxO validation, we further identify ET-insensitive populations and introduce a predictive therapeutic pipeline for actionable targeting. Together, our study provides a foundation for characterizing ET-insensitive populations at single-cell resolution, advances a translational framework for refining endocrine therapy, and supports the development of personalized strategies to overcome ET resistance in ER+ breast cancer.

## Methods

### Analysis of clinical samples from FELINE clinical trial

Serial single cell transcriptomes of tumor cells derived from patient biopsies collected during clinical trial FELINE <u>(#NCT02712723, registered on March 18, 2016 at ClinicalTrials.gov),</u> which enrolled postmenopausal women with ER-positive/HER2-negative breast cancer, were accessed through GEO under accession code GSE158724(11). Three tumor core biopsies were collected over the course of treatment: screening (Day 0), Cycle 1 Day 14 (Day 14), and end of trial (Day 180). To perform integration and comparative analysis of ET-sensitive and -insensitive cells, cell populations were extracted from each dataset and re-clustered as two independent datasets using integration analysis provided by Seurat package.

### Differential abundance analysis

To quantify changes in cell population abundance between conditions, we computed log2 fold-changes (enrichment scores) in relative abundance for each cluster. For the FELINE trial data, we compared Day 14 to Day 0 samples; for the PDxO models, we compared endocrine therapy (ET)-treated cells to Vehicle control. For each cluster X, the relative abundance was calculated as the proportion of cells in cluster Xdivided by the total number of cells in the corresponding condition. The log2 fold-change was then computed as:

FELINE trial:

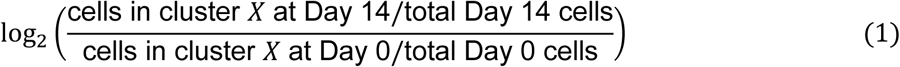

PDxO models:

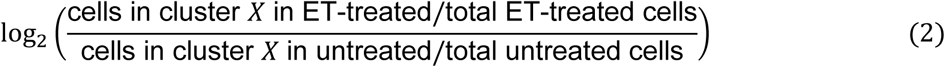

This approach normalizes for differences in total cell counts and enables direct comparison of cluster dynamics across conditions. Cell populations/clusters that were statistically significantly depleted with ET were considered to be ET-sensitive, while cell populations that were enriched or not affected by ET were considered to be ET-insensitive. For clusters with zero cells at either time point, a pseudocount of 1 was added to the zero-count group and to the corresponding non-zero group before calculating enrichment and log2 enrichment values to avoid undefined ratios. Statistical significance was determined using chi-square test based on the original observed cell counts.

### Simpson’s diversity index

Simpson’s diversity index (1 − D) was used to quantify heterogeneity of clusters across patients and PDxO models. For each cluster, the proportion of cells contributed by each patient or PDxO model was calculated, and Simpson’s diversity index was computed as:

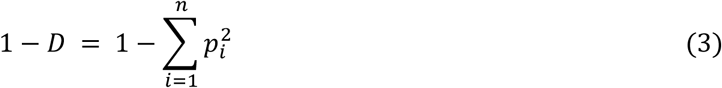

where *p_i_* is the proportion of cells within a cluster contributed by patient or PDxO model *i*. Higher index values indicate greater diversity, suggesting that the corresponding cell populations are more evenly distributed and shared across patients. The index was computed per patient and for aggregated groups (ET-sensitive vs. ET-insensitive). Diversity scores for each cluster were then stratified into ET-sensitive and ET-insensitive groups and visualized using dot plots.

### Functional enrichment analysis (FEA)

FEA was used to identify enrichment of gene signatures across the identified clusters(15). Signatures tested were derived from MSigDB v.7.4 or custom generated from the top 200 up-regulated genes identified using Seurat package(14,30,31). Prior to calculating signature scores, the data was normalized and scaled gene-wise. Then z-scored signatures were calculated for each cell separately. ROC analysis was used to estimate the accuracy of enrichment of a signature within a particular cluster. Area Under the Curve (AUC) > 0.6 was considered an enrichment. Significance of a signature enrichment across the clusters was estimated by the Wilcoxon rank-sum test (p < 0.01 was considered significant). For ER-associated gene signatures, composite ER scores were calculated as the mean of multiple ER-related signature scores per patient or cluster.

### Single cell molecular subtyping (SCSubtype)

Single-cell molecular subtyping was performed using the SCSubtype algorithm to assign intrinsic breast cancer subtypes, Luminal A, Luminal B, HER2-enriched, or Triple Negative, on a per-cell basis. This approach enables high-resolution classification of individual tumor cells based on their transcriptomic profiles(19).

### Gene signatures derived from ET-insensitive cell population in METABRIC dataset

Gene signatures were generated from the top 200 upregulated differentially expressed genes (DEGs) identified in ET-insensitive cell populations from patient-derived xenograft organoids (PDxOs) or patient tumors from the FELINE clinical trial. For each gene signature, survival and clinicopathological association analyses were performed using the METABRIC breast cancer cohort. Gene expression and clinical metadata of the METABRIC dataset were obtained from cBioPortal. For each signature, a composite score was calculated for each patient as the mean expression of signature genes. Patients were dichotomized into high- and low-expression groups using the median signature score. The median cutoff was selected a priori and applied uniformly across all signatures to provide a consistent threshold while avoiding outcome-driven cutoff optimization. Kaplan–Meier survival curves were generated using the dichotomized groups, and hazard ratios were estimated using univariable Cox proportional hazards models comparing the high- and low-expression groups. Associations with overall survival (OS) and relapse-free survival (RFS) were assessed using Kaplan–Meier analysis and log-rank tests implemented via the *survival* and *survminer* packages. Additional associations with clinical pathological features, including tumor grade, stage, nodal status, ER/PR/HER2 status, and PAM50 subtype, were evaluated using chi-square or Fisher’s exact tests, as appropriate. All statistical analyses were performed in R.

We used LASSO regression to identify genes that contribute most to the prediction of poor patient outcomes, including overall survival (OS) and relapse-free survival (RFS), within each gene signature (S. Table 11, 24). Gene expression matrices were restricted to genes present in each signature and matched to clinical data. For each signature, expression values were standardized, and missing values were imputed using the median expression of each gene. LASSO-Cox models were fitted using the glmnet R package with 10-fold cross-validation to determine the optimal penalty parameter (λ). The value of λ corresponding to the minimum cross-validated error (λmin) was used to select genes with non-zero coefficients, representing those contributing to the survival association. Analyses were performed separately for RFS and OS.

### Spatial and single cell RNA sequencing and data analysis

Spatial transcriptomic profiling was performed on FFPE-preserved primary tumors (UIC013 and UIC020) and matched PDX samples using the 10x Genomics Visium platform. Tissue sections were processed according to the manufacturer’s protocol, including permeabilization, reverse transcription, and library preparation. Library construction and sequencing was performed at the UIUC Roy J. Carver Biotechnology Center. The libraries were prepared with the Visium CytAssist Spatial Gene Expression kit from 10X Genomics. The libraries were pooled; quantitated by qPCR and sequenced on one 25B lane with 28×50nt reads on a NovaSeq X Plus with V1.0 sequencing kits. Raw spatial data were processed using Space Ranger (10x Genomics) to generate feature-barcode matrices mapping gene expression to specific spots on the Visium slide. All spatial data processing was conducted at the RRC Research Informatics Core at the University of Illinois Chicago (UIC).

To exclude potential murine contamination in PDX samples, each dataset was first aligned to the mouse and human reference genomes, and mouse-derived counts were removed from the final expression matrices. Spot-level annotation was performed using a reference breast cancer single-cell atlas(19) and Seurat’s label transfer method. Epithelial cell–enriched regions were identified based on marker gene expression profiles derived from the atlas.

Single-cell RNA sequencing was performed on dissociated PDxO cultures using the 10x Genomics Chromium Single Cell 3’ v3 platform. After 14 days of Vehicle, 4-hydroxy-tamoxifen (4OHT) (1µM), or Fulvestrant (ICI-182780) (1µM) treatment PDxOs were enzymatically dissociated into single-cell suspensions, filtered, and loaded into the Chromium controller. Libraries were prepared following the manufacturer’s protocol and sequenced on an S4 2×150nt lane in a NovaSeq 6000. Raw sequencing data were processed using 10x Genomics Cloud analysis for demultiplexing, alignment to the GRCh38 reference genome, and UMI counting.

### Data integration and analysis

Processed single-cell and spatial datasets were analyzed using Seurat (v4.3.0) in R. Quality control filters were applied to exclude cells with low gene counts, high mitochondrial content, or doublets. Normalization and scaling were performed using SCTransform(14). Dimensionality reduction was achieved via principal component analysis (PCA) and Uniform Manifold Approximation and Projection (UMAP). Clustering was performed using a shared nearest neighbor (SNN) modularity optimization algorithm.

Label transfer was used to map epithelial subpopulations/clusters identified in spatial datasets onto matched PDX and PDxO single-cell datasets, enabling assessment of subpopulation conservation across models. Differential expression analysis was conducted using the Wilcoxon rank-sum test, and pathway enrichment was assessed using MSigDB gene sets.

### Nanostring profiling on primary tumor and PDX models

NanoString Breast Cancer 360™ Biological Signatures analysis was conducted on primary tumors and matched PDX models for UIC013 and UIC020 at the University of Illinois Chicago (UIC). RNA was extracted from flash-frozen tumor tissues and quantified using a NanoDrop spectrophotometer. Samples were processed according to the manufacturer’s protocol for the Breast Cancer 360 panel, which includes 770 genes representing key biological pathways and molecular subtypes relevant to breast cancer. Hybridization, post-hybridization processing, and data acquisition were carried out using the nCounter® MAX Analysis System.

### DREEP analysis

To identify candidate drugs targeting ET-insensitive cell populations, we applied DREEP (Drug Estimation from single-cell Expression Profiles) analysis(20). For each ET-insensitive cluster identified through scRNA-seq, the top 200 differentially expressed genes (DEGs) were selected and used as input for DREEP. This analysis includes precomputed sensitivity biomarkers for each of the 450 drugs, obtained from the publicly available The Cancer Therapeutics Response Portal v2(32). A positive score indicates drug resistance, while a negative score indicates sensitivity. The result of the DREEP analysis is a list of sensitivity scores for 450 tested drugs with associated p-values generated for each cell population. DREEP was run at the cell population level to predict compounds with selective efficacy against each cluster.

Each DREEP compound was assigned to a biological pathway based on its known molecular target. For each pathway, gene signatures were curated from MSigDB (S. Table 29, 30). Pathway activity was calculated for each cluster using gene signature scoring implemented in Seurat (e.g., AddModuleScore) based on normalized gene expression data. To identify the connection between DREEP-predicted drug sensitivities to transcriptional pathway activity across ET-insensitive clusters, we calculated correlations between DREEP drug sensitivity scores (DSS) and pathway gene signature scores across clusters (n = 9) for each compound. Pearson correlation was used as the primary measure due to the limited number of clusters, with Spearman correlation calculated as a complementary non-parametric approach. Associated p-values were obtained using two-sided tests and adjusted for multiple hypothesis testing using the Benjamini– Hochberg method. To assess overall trends, we analyzed the distribution of correlation coefficients across all drug–pathway pairs, testing whether the proportion of negative correlations differed from the expected null proportion (0.5) using a binomial test, and whether mean correlation coefficients per pathway differed from zero using a one-sample t-test.

### Adaptation of mixed model with patient-specific random intercepts for PDxO signatures in FELINE

To assess temporal changes in gene signature scores at single-cell resolution, we adapted a hierarchical modeling approach from Griffiths et al. to analyze PDxO-derived signature scores(11). Briefly, average gene signature scores for each cell were calculated using the AddModuleScore() function in the Seurat R package. We next implemented a linear mixed effects framework that leveraged all available cells while accounting for patient-level variability, enabling robust inference of treatment response dynamics across molecular signatures. Single-cell signature scores were computed using predefined gene sets. Each cell was annotated with treatment arm, clinical response category (Responsive vs. Non-responsive), and time since treatment initiation. To improve model stability, time was rescaled by a factor of 100. For each treatment arm and gene signature, we modeled single-cell pathway activity using a linear mixed-effects framework:

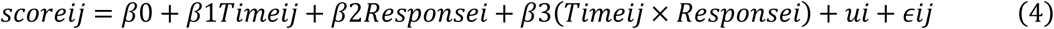

where:

score_ij_ is the signature score for cell j from patient i
Time_ij_ is the rescaled time point (e.g., days since treatment initiation)
Response_i_ is a binary indicator of treatment response (Responsive vs. Non-responsive)
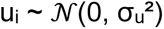 is a patient-specific random intercept
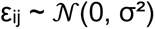 is the residual error

The interaction estimate (β_3_) quantifies the difference in the temporal trajectory of signature activity between clinically responsive and non-responsive tumors. Positive values indicate greater increases in signature activity over time in non-responsive tumors relative to responsive tumors, whereas negative values indicate the opposite pattern.

Models were fit using the lmer() function from the *lme4* R package with restricted maximum likelihood (REML) estimation and the bobyqa optimizer. Likelihood ratio tests were used to evaluate the significance of time effects and response-specific trends. For each signature, we performed likelihood ratio tests comparing: 1. the full model vs. a reduced model excluding time (to test for any temporal change), and 2. the full model vs. a model assuming equal slopes across response groups (to test for differential dynamics). For each signature, Supplementary Table 23 reports the number of patients included in the analysis (N_patients), the total number of cells contributing to the model (N_cells_total), the estimated response-by-time interaction coefficient (Interaction_estimate), its corresponding Wald p-value (Interaction_p), and the p-values from the two likelihood ratio tests (Lrt_noTime_p and Lrt_equalChange_p). The p-values displayed in Figure 7D–E correspond to the response-by-time interaction (Interaction_p). Predicted trajectories were visualized using ggplot2, with separate curves for Responsive and Non-responsive groups.

The longitudinal mixed-effects analysis included all patients with cells available for gene signature scoring (N = 11). The analysis incorporated all available samples collected at Day 0, Day 14, and Day 180; however, not all patients contributed samples at every timepoint. This differs from the integrated clustering analyses presented in Figures 1 and 2, which were restricted to nine patients with sufficient numbers of cells at both Day 0 and Day 14 to enable reliable patient-specific clustering. Because gene signature scoring does not require de novo clustering, two additional patients were included in the longitudinal analysis.

To evaluate the robustness of the cell-level analysis and address potential pseudoreplication, we additionally performed a patient-level sensitivity analysis by averaging signature scores within each patient-timepoint sample. Patient-level mean signature scores were analyzed using linear mixed-effects models including fixed effects for clinical response, categorical timepoint (Day 0, Day 14, and Day 180), and their interaction, with patient included as a random intercept. Estimated marginal means and statistical results from this sensitivity analysis are provided in Supplementary Table 24.

### PDX development

Fresh tumor tissue was obtained from patients undergoing primary breast surgery at the University of Illinois Hospital, Chicago, IL, USA in accordance with the Declaration of Helsinki. The study was reviewed by the Institutional Review Board of the University of Illinois at Chicago (IRB #1, protocol number: 2014-1205) and was determined not to involve human subjects research, as all data and specimens were de-identified; therefore, informed consent was not required in accordance with 45 CFR 46.102. Tumor fragments were implanted into the cleared mammary fat pads of immunodeficient NSG (NOD scid gamma) mice under anesthesia with ketamine (100 mg/kg) and xylazine (10 mg/kg), administered intraperitoneally(18). Mice were maintained in University of Illinois at Chicago Biologic Resources Laboratory (BRL)-managed animal facilities and housed in sterile static microisolator cages on autoclaved corncob bedding, with irradiated food and autoclaved water provided ad libitum. Animals were maintained under a 14-hour light/10-hour dark cycle, and cages were changed at least weekly in a biosafety cabinet. Randomization and blinding were not applicable to this study, as no interventional or comparative experimental procedures were performed. All animal procedures were approved by the University of Illinois at Chicago Office of Animal Care and Institutional Biosafety (OACIB), Animal Care Committee (ACC Protocol #23-010) and were conducted in accordance with institutional guidelines and the Guide for the Care and Use of Laboratory Animals. Mice were euthanized using carbon dioxide (CO_2_) inhalation at a displacement rate of approximately 30% of the chamber volume per minute, followed by cervical dislocation as a secondary physical method to ensure death, in accordance with AVMA guidelines.

### PDxO development

PDxO cultures were developed from freshly harvested PDX tumors and cultured following established protocol(18). Briefly, tumors were minced and enzymatically digested using collagenase/hyaluronidase followed by mechanical dissociation. The resulting single-cell suspensions were embedded in Matrigel and cultured in organoid medium optimized for ER+ breast cancer(18). Organoids were passaged every 10–14 days and maintained under standard conditions (37°C, 5% CO_2_). To eliminate murine contamination, organoid cultures were either subjected to differential centrifugation following dispase digestion or sorted by FACS using species-specific surface markers. Post-sorting, human cells were aggregated overnight in ultra-low attachment plates and embedded in Matrigel the following day; minimal mouse content was confirmed by RT–qPCR using mouse GAPDH primers.

### Immunohistochemistry for ERα on PDX Models

Immunohistochemistry (IHC) was performed to assess estrogen receptor alpha (ERα) expression in formalin-fixed, paraffin-embedded (FFPE) PDX tumor sections. Tissue sections were deparaffinized, rehydrated, and subjected to antigen retrieval using citrate buffer (pH 6.0) at 95°C for 20 minutes. Endogenous peroxidase activity was blocked with 3% hydrogen peroxide, followed by incubation with a primary antibody against ERα (clone SP1, Catalog No. RM-9101-S1, Fisher Scientific) overnight at 4°C. Detection was carried out using a biotin-streptavidin HRP system and DAB chromogen. Samples processing and imaging were conducted at UIC RRC Histology Core.

### Immunofluorescence for ERα on PDxO models

PDxO cultures were dissociated to single cells by enzymatic digestion. The resulting cell suspension was seeded onto glass coverslips in estrogen-deprived medium (10% CD-FBS Phenol red free) and allowed to adhere for 24 hours. Cells were then fixed with 4% paraformaldehyde, permeabilized, and blocked with casein for 1 hour at room temperature. ERα staining was performed using an undiluted primary antibody (Abcam #ab166600) incubated overnight at 4°C. After washing with 1× TBS, cells were incubated with Alexa Fluor 495 secondary antibody for 1 hour at room temperature. Coverslips were washed with x1 TBS and mounted using ProLong™ Gold Antifade Mountant with DAPI. Samples were imaged using a Leica DMi8 fluorescence microscope.

### Proliferation assay on PDxO models

PDxO cultures were harvested by digesting Matrigel domes with dispase (50 U/ml) supplemented with 200 µl FBS and 1 µl ROCK inhibitor. Domes were scraped into the dispase mixture and mechanically disrupted to release organoids. After washing, organoids were resuspended in fresh Matrigel at a final concentration of 5,000 organoids/ml. A 20 µl Matrigel/organoid suspension was plated per well in a 48-well plate to form domes. Organoids were cultured in PDxO medium with chosen compounds from DREEP analysis. Media were refreshed every 3-4 days. Organoid growth was analyzed on 14th day using the Celigo image cytometer with the colony analysis module, which quantified individual organoid area (µm^2^). Dose-response assays were performed in two or three independent biological experiments, depending on the compound. For each biological replicate, the area under curve (AUC) was calculated using GraphPad Prism. AUC values were compared among PDxO models using one-way ANOVA followed by Dunnett’s multiple-comparisons test (S. Table X).

### EdU staining in PDxO models

PDxOs were dissociated, seeded in 6-well plate (n=3 per treatment group), and treated with vehicle control (Veh), 4-hydroxytamoxifen (4OHT, 1 µM), or fulvestrant (ICI, 1 µM) for 2 weeks, with media refreshed every 3–4 days. Following treatment, organoids were incubated with 10 µM 5-ethynyl-2′-deoxyuridine (EdU) for 4 hours. Organoids were then enzymatically dissociated into single-cell suspensions, fixed in 4% paraformaldehyde, and permeabilized prior to detection using the Click-iT™ EdU Alexa Fluor 647 assay according to the manufacturer’s protocol. EdU incorporation, reflecting S-phase entry, was detected via copper-catalyzed click chemistry. Nuclei were counterstained with DAPI, and samples were analyzed by flow cytometry with CytoFLEX S Flow Cytometer. Data were processed using FlowJo v11.1.

### Cell viability assay

MCF-7 cells and their endocrine therapy–resistant derivatives (TAMR, FULR, and HER2) (21,22) were seeded in triplicate at 2,500 cells per well in 96-well plates using their respective standard growth media. Cells were treated with compounds predicted by DREEP analysis in a dose– response format. On day 5, cells were stained with Hoechst33,4322 (LifeTechnologies, Carlsbad, CA, USA) and propidiumiodide (PI) (final concentration 1µg/mL) for 30min at 37C to quantify total cell number and percent cell death. Plates were scanned using the Celigo image cytometer, and data were analyzed using the dead/total module to retrieve total cell counts and viability metrics. Additionally, growth rate (GR) metrics were applied to normalize drug response across cell lines with differing proliferation rates(33).

### Clonogenic assay of cell lines

MCF-7 cells were seeded as single cells at 300 cells per well in 24-well plates under standard growth conditions, in triplicate. The following day, cells were treated with various doses of compounds predicted by DREEP analysis, with or without 1 µM 4-hydroxytamoxifen (4OHT). Treatments were refreshed every 3–4 days, and cells were cultured for 14 days to allow colony formation. On day 14, cells were stained with Hoechst and imaged using the Celigo image cytometer. Colony confluence was quantified for each condition using the Celigo analysis software.

## Data availability

ScRNA-seq data from the FELINE clinical trial can be found Gene Expression Omnibus (GEO) platform under the accession code GSE158724. Raw and demultiplexed single-cell RNA sequencing data are available through GEO under accession code GSE310968 and spatial RNA-seq data from PDxO models are under accession code GSE310362.

## Code availability

Code available on GitHub: https://github.com/sesemina/ProfilingETinsensitivePopulations.

## Authors’ Contributions

**S. E. Semina:** contributed to the conception and experimental design of the work, bioinformatics analysis, manuscript writing. **R.J. Huggins, H. Zhao, K. Yanagihara** and **M. Sheinin:** PDX and PDxO development, manuscript writing. **V. Macias and A. Balla:** pathological assessment of tumor specimen, **F. Alani**: aided with experiments setup. **L. Feferman:** contributed to bioinformatics analysis. **D. A. Tonetti, G. L. Greene, J. Frasor:** contributed to the conception design, models development, data analysis and manuscript writing. **K. F. Hoskins:** contributed to clinical specimen acquisition, clinical data collection, manuscript writing, and the project design. **J. Coloff:** contributed to conceptualization, supervision, writing manuscript draft, review and editing, and data analysis and interpretation.

## Acknowledgements

This work was supported by the National Institutes of Health (NIH) under award numbers R01 CA200669 and R21 CA276820. The funding sources had no involvement in the study design, data collection, analysis, or manuscript preparation.

The authors thank **Dr. Rachel Schiff** (Baylor College of Medicine, Houston) for generously providing endocrine therapy-resistant cell lines. We also thank **Dr. Alana Welm** for providing the HCI003 and HCI017 PDxO models. We are grateful to **Dr. Zarema Arbieva**, Director of the Genome Research Core, and **Dr. Mark Maienschein-Cline**, Director of the Research Informatics Core at the University of Illinois Chicago (UIC) Research Resources Center and Director of the Cancer Bioinformatics Shared Resource (CBSR) in the University of Illinois Cancer center, for their invaluable assistance with spatial and single-cell RNA-sequencing data acquisition and analysis. The Research Informatics Core is supported in part by NCATS through Grant UM1TR005438. We further acknowledge **Dr. Alvaro G. Hernandez**, Director of DNA Services, and **Dr. Mayandi Sivaguru**, Director of the Cytometry and Microscopy to Omics Facility, at the Roy J. Carver Biotechnology Center, University of Illinois Urbana–Champaign (UIUC), for their support in spatial and single-cell transcriptomics data generation. We thank **Dr. Maria Sverdlov**, Director of the Histology and Tissue Imaging Core and the UI Health Biorepository, for her assistance with spatial RNA-seq data acquisition. We also thank **Dr. Balaji Ganesh**, Director of the Flow Cytometry Core at UIC, for the assistance with EdU assay. Finally, we greatly appreciate **Aanshi Vashi** and **Godwin K. Sarpey** for their provided laboratory support and assistance with experiments.

