## Supplemental Figures for "Integrative pipeline to profile and target endocrine therapy-insensitive cell populations in ER+ breast cancer"

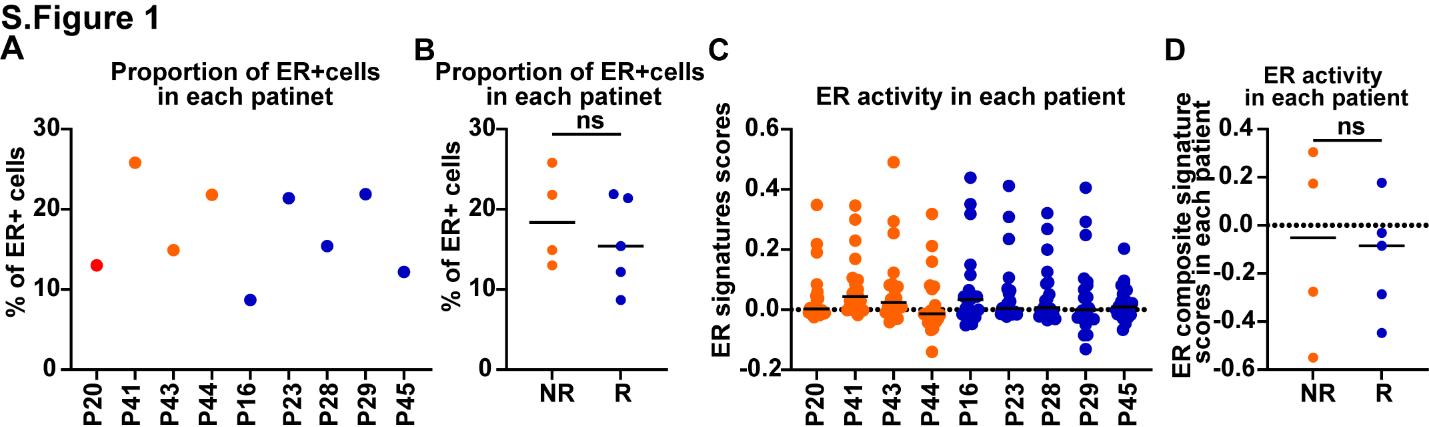

**Supplementary Figure 1. ER expression and activity across patients.** **(A)** Proportion of ER+ cells in individual patients. **(B)** Comparison of ER+ cell proportions between non-responsive (NR) and responsive (R) patients. **(C)** ER signaling activity scores per patient. Each point represents a score for individual gene signature per patient. ER-associated gene signatures were derived from MSigDB. **(D)** Composite ER scores (mean of ER-associated signature scores per patient) grouped by clinical outcome. Each point represents an individual patient; horizontal lines indicate group means. For **(B)** and **(D)** statistical comparisons were performed using unpaired t test, ns = not significant.

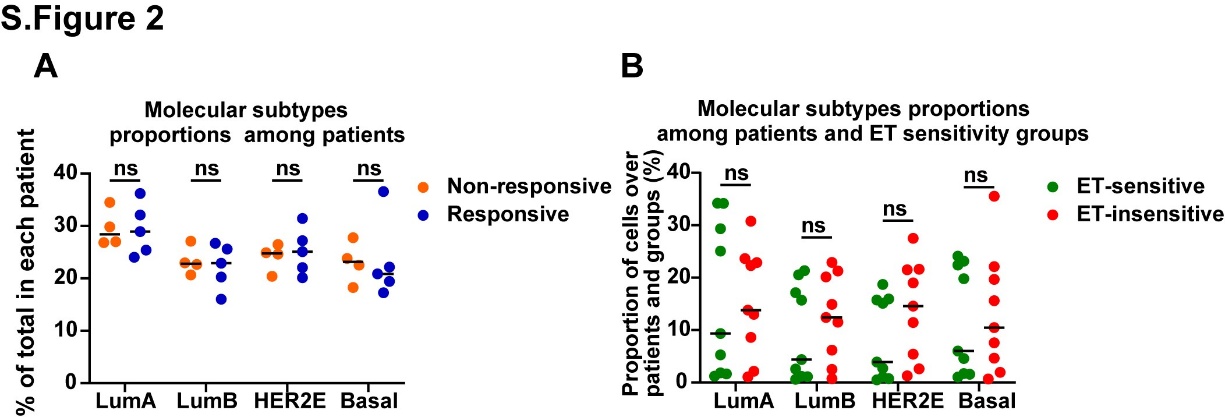

**Supplementary Figure 2. Molecular subtype distribution across patients and endocrine therapy response groups. (A)** Proportions of molecular subtypes (LumA, LumB, HER2E, Basal) across individual patients, stratified by clinical response. **(B)** Proportions of molecular subtypes across patients grouped by ET sensitivity. Each point represents an individual patient; horizontal lines indicate group means. Differences in molecular subtype proportions were assessed using two-way ANOVA, ns = not significant

**Supplementary Figure 3.** **Molecular fidelity of UIC013 and UIC020 PDX models.** **(A–D)** Wheel plot shows results of NanoString profiling of primary tumor and associated PDX. (TIS = Tumor Inflammation Signature) **(E–F)** ERα expression was assessed by IHC in UIC013 and UIC020 PDX tumors. Scale bar = 50 µm.

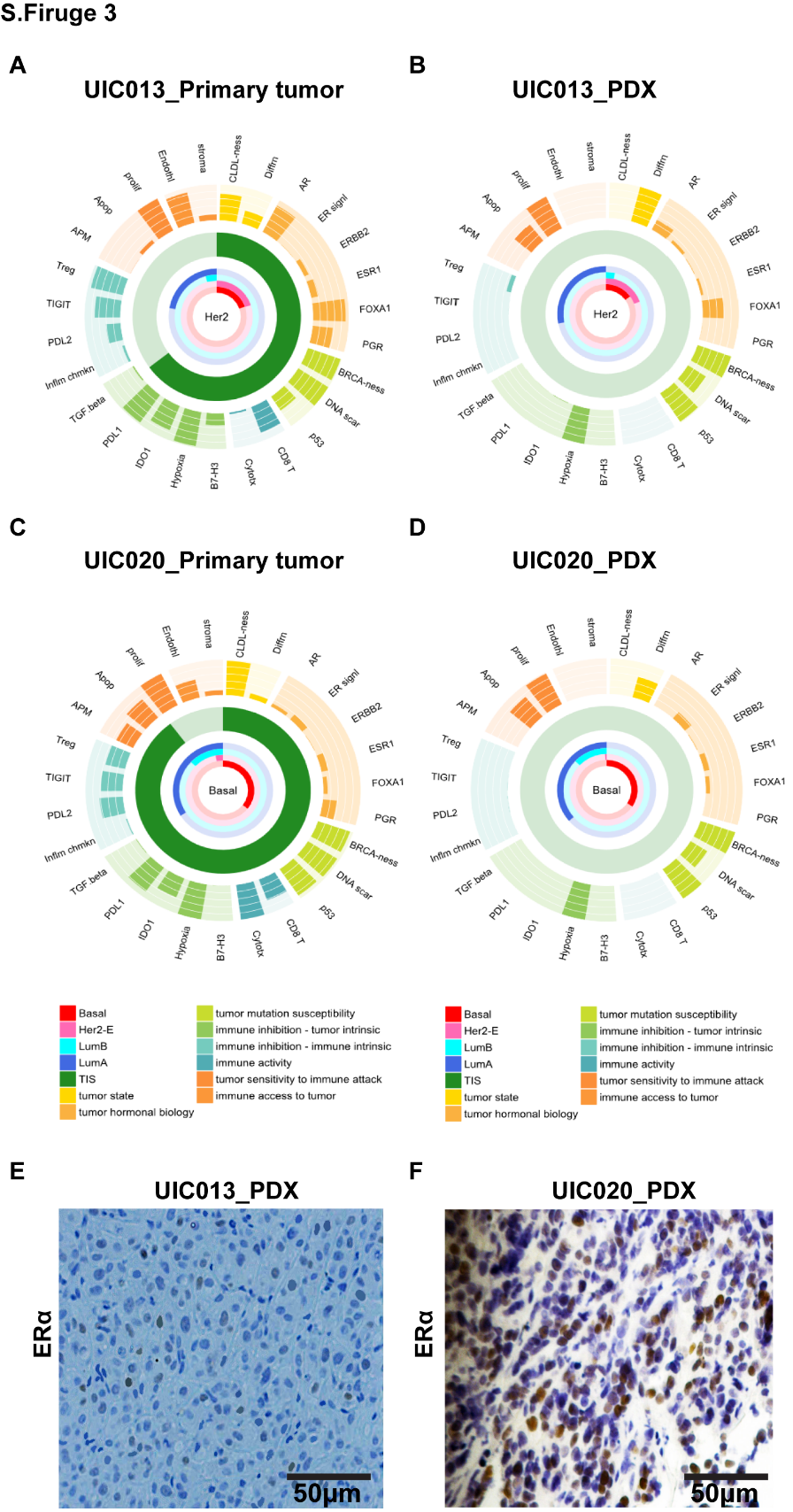

**Supplementary Figure 4.** **Newly developed UIC013 PDxO model retains molecular and cellular heterogeneity of ER+ breast tumors.** **(A–C)** Schematic representation of matched primary tumor, PDX, and PDxO models for UIC013 analysis. **(A)** Schematic overview of models’ development. **(B)** Visual representation of each UIC013 model. Hematoxylin and eosin (H&E) staining was used to visualize primary and PDX tumors, and brightfield microscopy was used to visualize PDxOs. Morphological annotations of the primary tumor regions were labeled: black—tumor, red—fibrosis, blue—necrosis, yellow—chronic inflammation. **(C)** Spatial clustering of primary tumor and PDX samples aligned to H&E-stained sections. UMAP plots show clustering of PDxO cells. **(D)** UMAP plot of breast cancer atlas with cell types labeled. The label transfer analysis by Seurat package was used to generate transcriptional profile of cancer epithelial cells. **(E)** Bar plot displays cell type distribution across spots in spatial RNA-seq of UIC013 primary tumor and PDX. **(F)** UMAP plot shows clustering of cancer epithelial spots from UIC013 primary tumor. **(G)** Bar plot shows the distribution of epithelial clusters from the UIC013 primary tumor across PDX and PDxO models. Label transfer analysis was used to assess conservation of epithelial subpopulations across models.

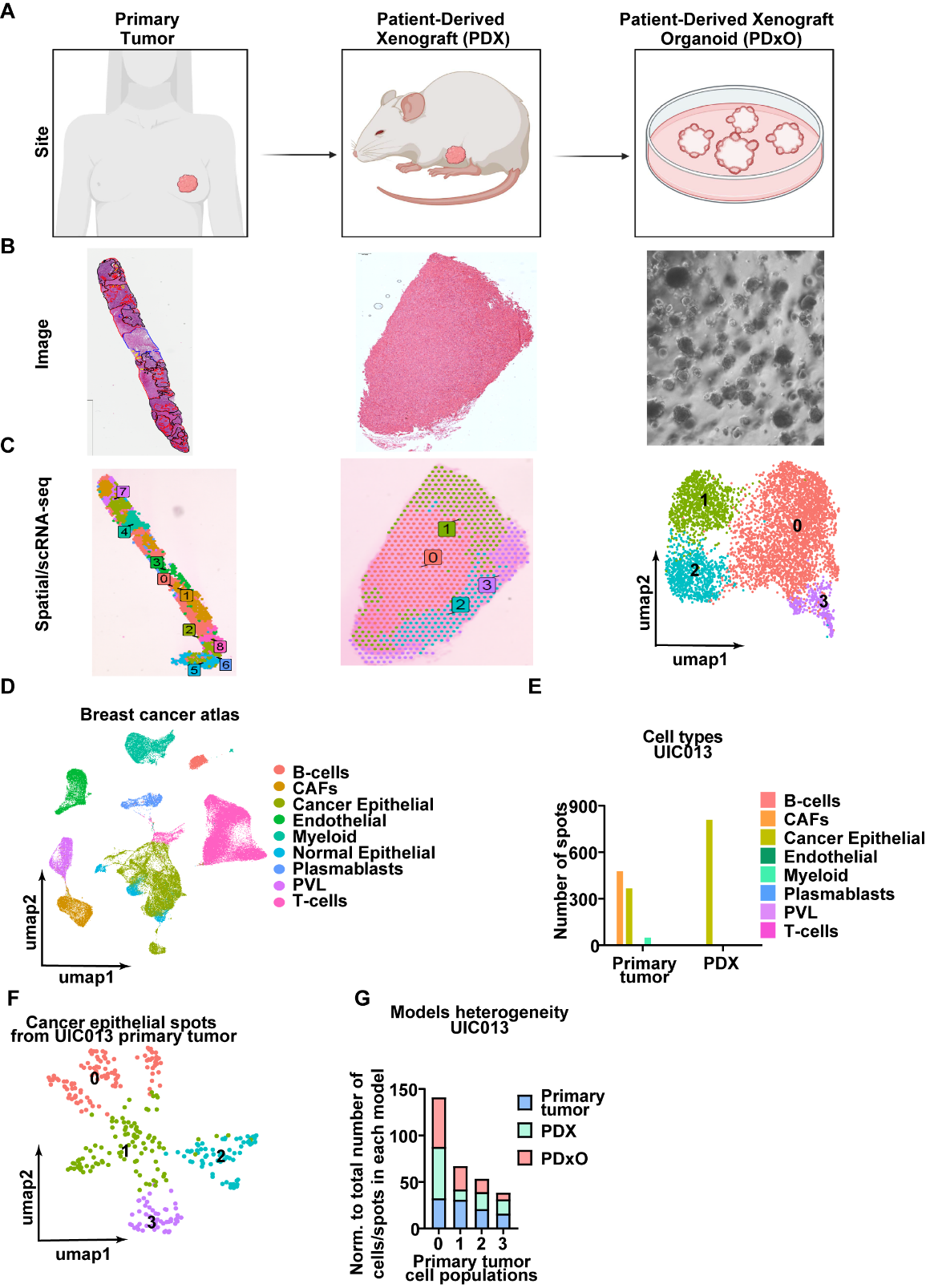

**Supplementary Figure 5.** **ER expression and functionality in new PDxO models UIC013 and UIC020.** **(A)** Representative immunofluorescence images showing ERα (red), DAPI (blue), and merged signals across PDxO models (HCI003, HCI017, UIC013, UIC020). **(B)** Quantification of ER+ cells across models. **(C–D)** UIC013 and UIC020 PDxOs were treated with fulvestrant (ICI-182,780), 4-hydroxytamoxifen (4OHT), or vehicle control (Veh) for 14 days. Data represents fold change in organoid area on day 14 of treatment as determined by Celigo image cytometer. **(E, G)** EdU-based cell cycle analysis showing the percentage of cells in G1, S, and G2/M phases following treatment after 14 days of treatment in each model. **(F, H)** Representative flow cytometry plots of EdU incorporation and DNA content (DAPI) for indicated treatments. Data are presented as mean ± SD; statistical significance indicated as ns (not significant), **p < 0.01, ****p < 0.0001.

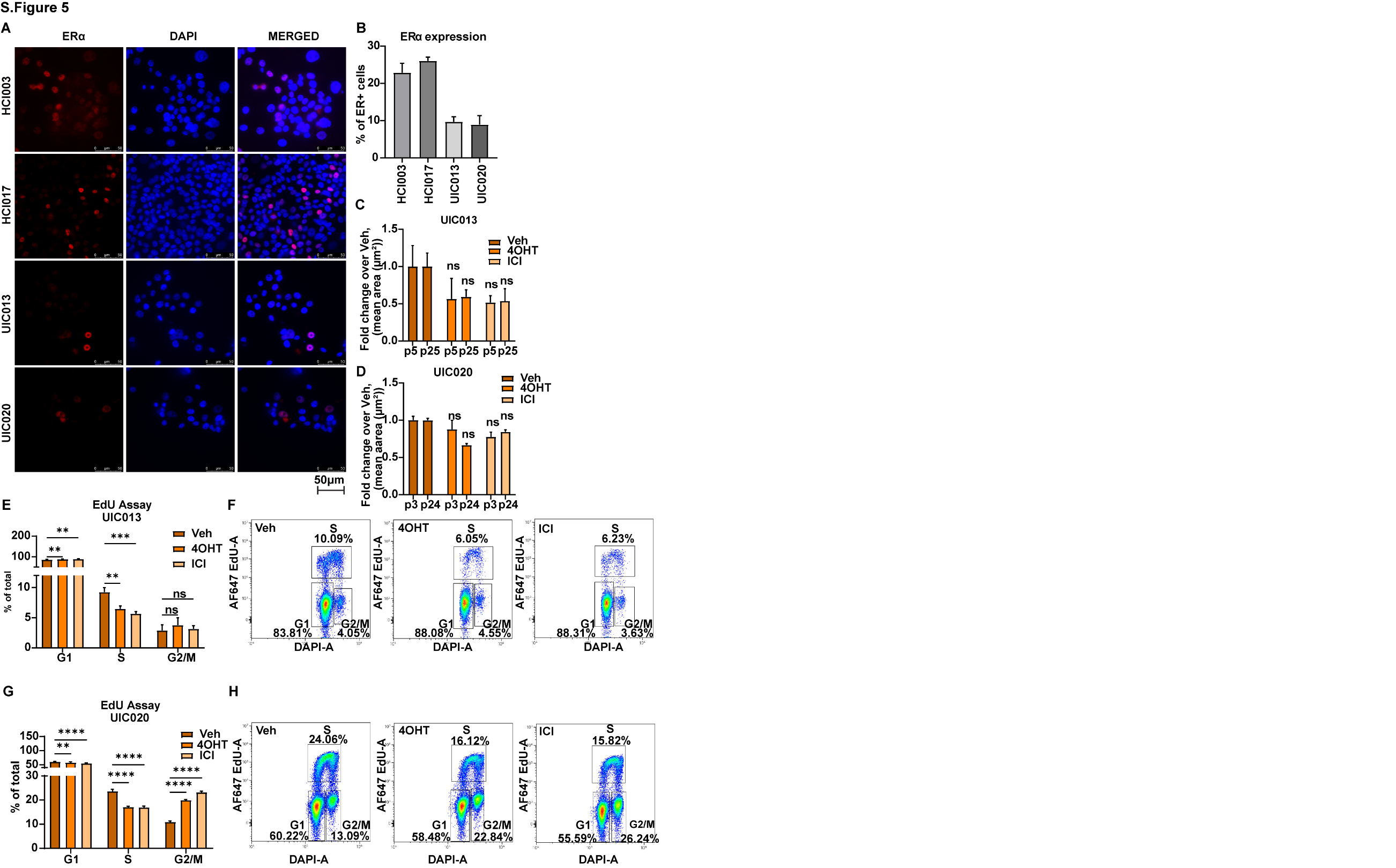

**Supplementary Figure 6. ER expression and activity at single cell resolution.** **(A–D)** UMAP plots of scRNA-seq data from each PDxOs. **(E)** Proportion of ER+ cells in each cluster identified in each model. **(F)** Proportion of ESR1+ cells in each cluster for each model is shown in bar plot. **(G)** Functional enrichment analysis (FEA) of ER gene signatures from MSigDB was performed on each cluster across all models. AUC values are shown in a heatmap, and p-values are presented in S. Table 15.

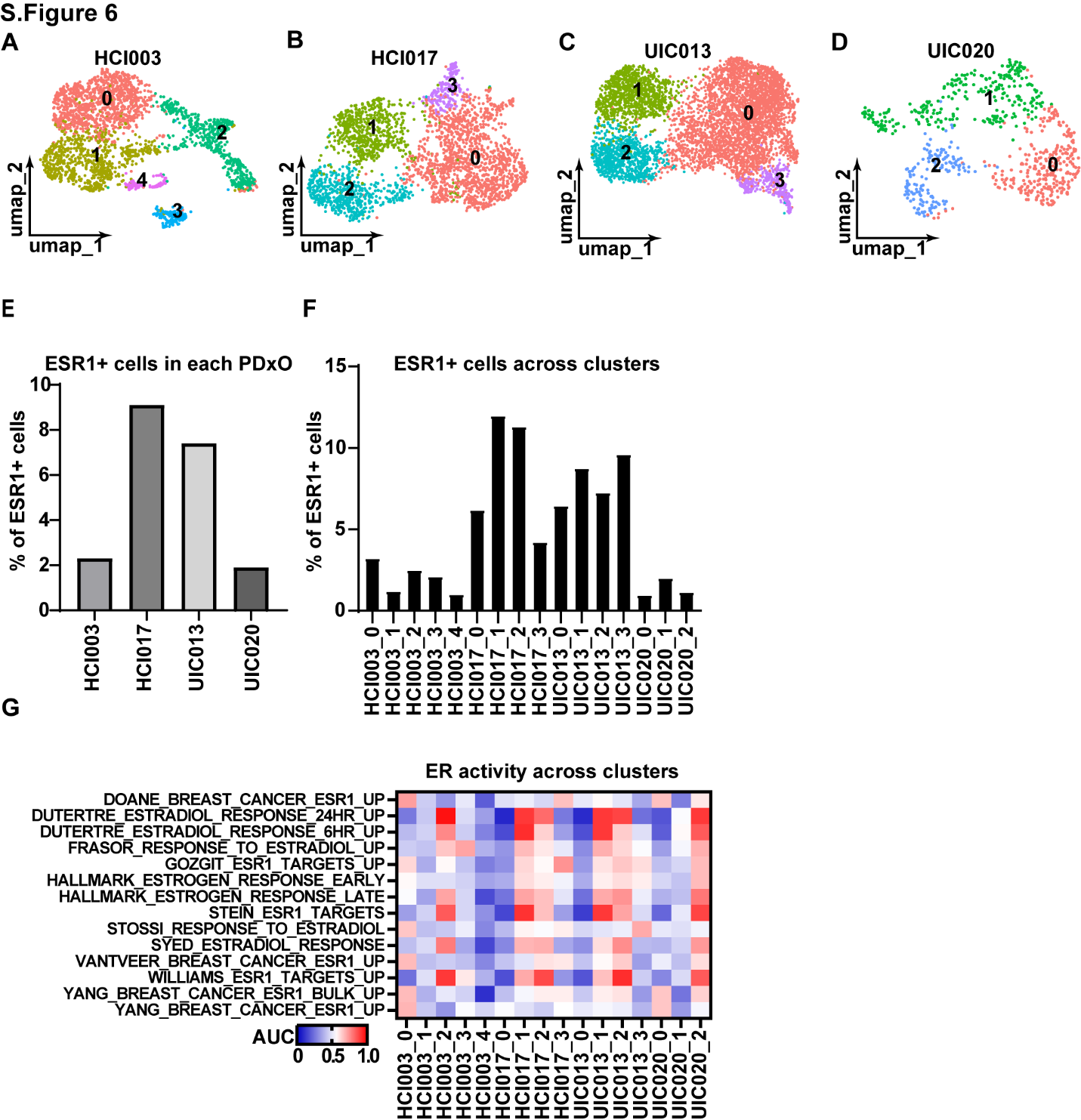

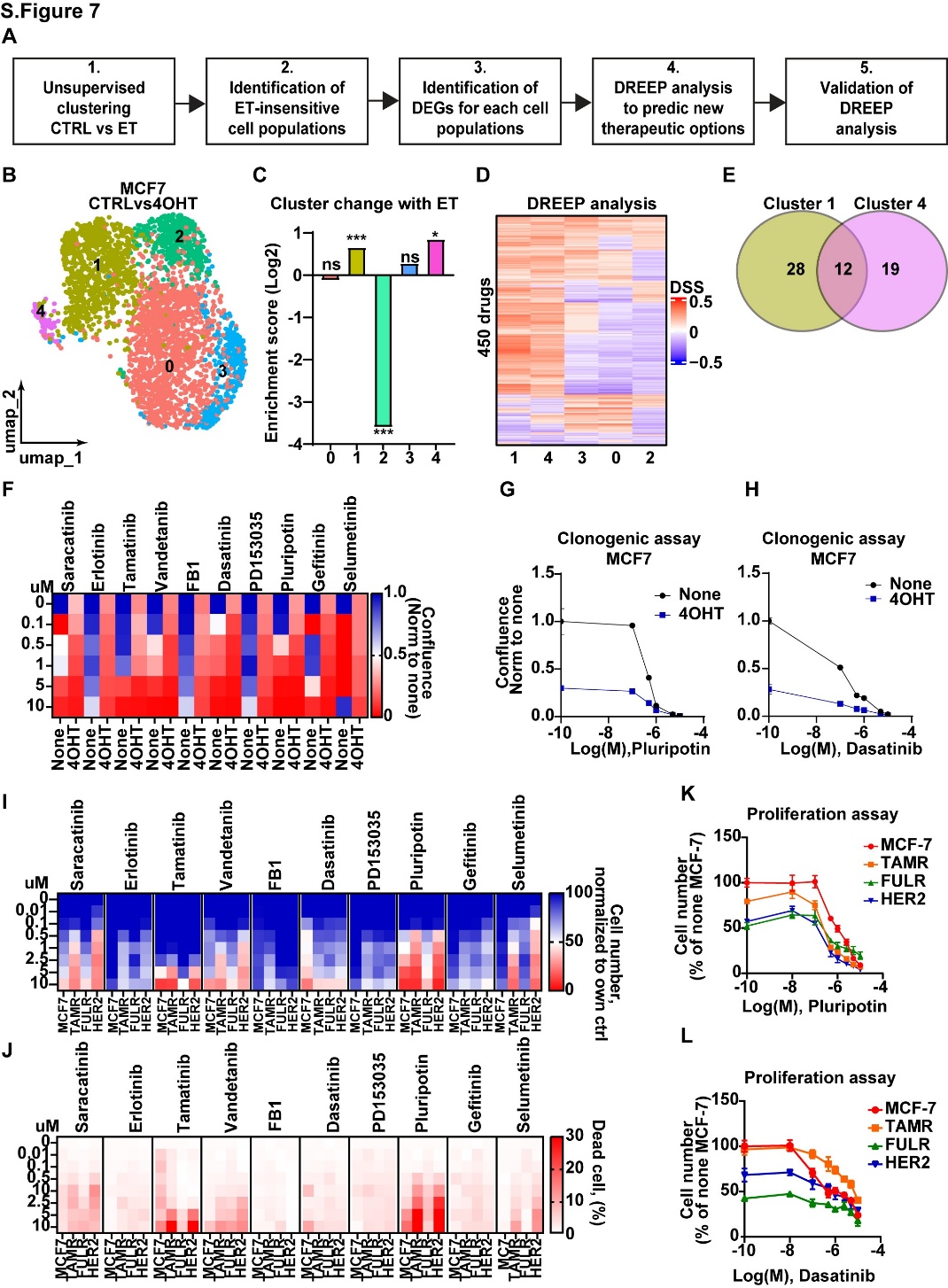

**Supplementary Figure 7. Predictive therapeutic pipeline identifies drugs targeting ET-insensitive populations from MCF7 cell line**. **(A)** Schematic representation of the new therapeutic pipeline integrating clustering, DEGs selection, and DREEP analysis. **(B)** UMAP plot shows clustering of single cells from MCF7 cells treated with 1 µM 4OHT or vehicle (CTRL) for 14 days. **(C)** Differential abundance analysis displays enrichment score and ET-sensitive clusters or -insensitive clusters. *P < 0.05, ***P < 0.001, ns = not significant.  **(D)** Heatmap depicts drug sensitivity scores (DDS) for 450 drugs tested for across clusters identified in panels A and B. Negative values indicate sensitivity, positive values indicate resistance. **(E–H)** Drug screening of 10 out of 12 compounds identified in **(E)** using clonogenic assays in MCF7 cells. Cell confluence, reflecting clonogenic growth of ET-insensitive cells, is presented as normalized to untreated controls in a heatmap **(F).** Columns indicate vehicle control (“None”) or treatment with a fixed dose of 4-hydroxytamoxifen (4OHT; 1 μM), while rows represent increasing concentrations of each compound. The 0 μM row corresponds to single-agent conditions, where “None” represents vehicle control and “4OHT” represents 4OHT alone. The dose-response curves for the two most effective drugs, Pluripotin **(G)** and Dasatinib **(H)**. **(I-L)** Drug screen of the same 10 drugs using viability assays in MCF7 and its ET-resistant derivatives. Total cell number **(I)** and proportion of dead cells **(J)** are shown in heatmaps, and as dose-response curves for Pluripotin **(K)** and Dasatinib **(L)**.

**Supplementary Figure 8.** **Growth rate–corrected drug response in ET-resistant cell lines.** **(A)** Schematic representation of growth rate (GR) metrics interpretation. **(B)** Dose-response curve for MCF7 and its ET-resistant derivatives treated with Pluripotin, analyzed using the GR algorithm. **(C)** Heatmap shows drug screen data from S. Figure 7I re-analyzed using GR metrics.

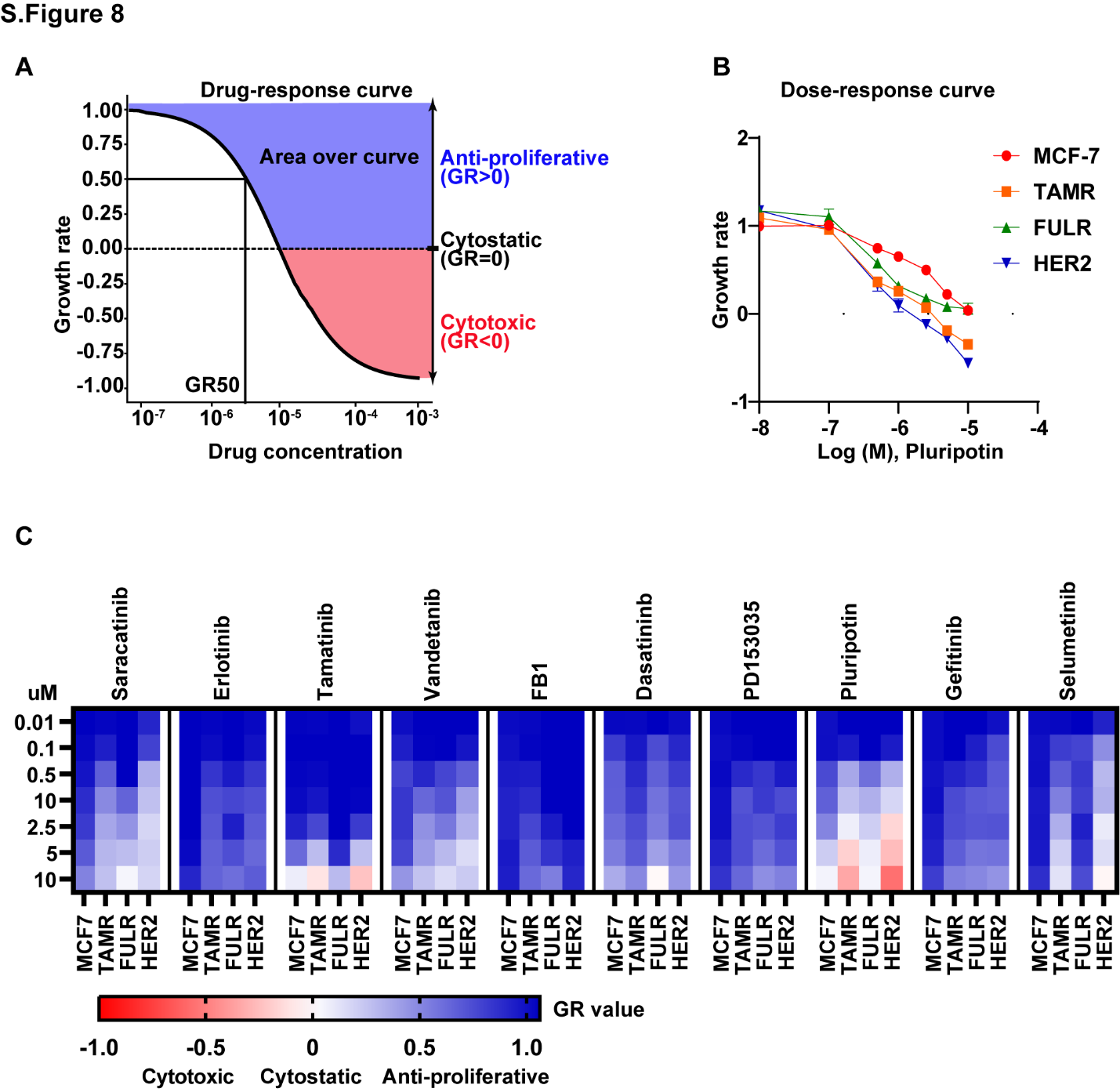

Supplementary Table Legends

Supplementary Table 1. Clusters enrichment with endocrine therapy in patient tumors from FELINE clinical trial.

Supplementary Table 2. FEA for ER associated gene signatures in clusters derived from patient tumors of FELINE clinical trial.

Supplementary Table 3. Statistical analysis of ER associated gene signatures across sensitivity and clinical response groups derived from patient tumors of FELINE clinical trial.

Supplementary Table 4. Statistical analysis of ER associated gene signatures across at the patient tumor level in FELINE clinical trial.

Supplementary Table 5. Distribution of ET-sensitive cells across clusters and patients in the FELINE trial.

Supplementary Table 6. Distribution of ET-insensitive cells across clusters and patients in the FELINE trial.

Supplementary Table 7. FEA for Reactome gene signature for ET-sensitive cell populations derived from FELINE clinical trial.

Supplementary Table 8. FEA for Reactome gene signature for ET-insensitive cell populations derived from FELINE clinical trial.

Supplementary Table 9. Top 200 DEGs of ET-insensitive cell populations derived from FELINE clinical trial.

Supplementary Table 10. Prognostic value of gene signatures derived from ET-sensitive and -insensitive cell populations of FELINE trial in METABRIC Breast Cancer _Survival.

Supplementary Table 11. Prognostic value of gene signatures derived from ET-insensitive cell populations of FELINE trial in METABRIC Breast Cancer dataset_Clinical Features.

Supplementary Table 12. LASSO regression on predictive gene signatures of ET-insensitive clusters derived from FELINE clinical trial.

Supplementary Table 13. Summary table for LASSO regression analysis on predictive gene signatures of ET-insensitive clusters derived from FELINE clinical trial.

Supplementary Table 14. Clinical characteristics of patients whose tumors were used to establish UIC and HCI patient-derived organoid (PDxO) models.

Supplementary Table 15. FEA for ER associated gene signatures in clusters derived from PDxO.

Supplementary Table 16. Clusters enrichment with endocrine therapy in PDXO models.

Supplementary Table 17. FEA for ER associated gene signatures in clusters derived from PDxOs treated with ET.

Supplementary Table 18. Statistical analysis of ER associated gene signatures across ET-sensitive and ET-insensitive cell populations derived from PDxO models.

Supplementary Table 19. Cluster composition of ET-sensitive and ET-insensitive cells from PDXO.

Supplementary Table 20. FEA for Reactome gene signature for ET-sensitive cell populations derived from PDxO models.

Supplementary Table 21. FEA for Reactome gene signature for ET-insensitive cell populations derived from PDxO models.

Supplementary Table 22. Top 200 DEGs of ET-insensitive cell populations derived from PDxO models.

Supplementary Table 23. Temporal changes in PDxO ET-insensitive gene signature scores at single-cell resolution in samples from FELINE clinical trial.

Supplementary Table 24. Patient-level sensitivity analysis of longitudinal PDxO-derived ET-insensitive gene signatures in the FELINE cohort.

Supplementary Table 25. Prognostic value of gene signatures derived from ET-insensitive cell populations of PDxO in METABRIC Breast Cancer _Survival.

Supplementary Table 26. Prognostic value of gene signatures derived from PDxO ET-insensitive cell populations in METABRIC Breast Cancer dataset_Clinical Features.

Supplementary Table 27. LASSO regression on predictive gene signatures of ET-insensitive clusters derived from FELINE clinical trial.

Supplementary Table 28. Summary table for LASSO regression analysis on predictive gene signatures of ET-insensitive clusters derived from FELINE clinical trial.

Supplementary Table 29. DREEP analysis for MCF7 cell populations treated with endocrine therapy.

Supplementary Table 30. DREEP analysis for PDxO ET-insensitive cell populations.

Supplementary Table 31. Statistical analysis of dose-response assays validating DREEP-predicted compounds in PDxO models.

Supplementary Table 32. Description of DREEP compounds and assigned pathways.

Supplementary Table 33. DREEP compounds targets and pathways correlation for ET-insensitive cell populations derived from PDxOs.

Supplementary Table 34. DREEP analysis for FELINE ET-insensitive cell populations.

Supplementary Table 35. DREEP analysis compound status summary for FELINE ET-insensitive cell populations.
